# Defect patterns on the curved surface of fish retinae suggest mechanism of cone mosaic formation

**DOI:** 10.1101/806679

**Authors:** Hayden Nunley, Mikiko Nagashima, Kamirah Martin, Alcides Lorenzo Gonzalez, Sachihiro C. Suzuki, Declan Norton, Rachel O. L. Wong, Pamela A. Raymond, David K. Lubensky

## Abstract

The outer epithelial layer of zebrafish retinae contains a crystalline array of cone photoreceptors, called the cone mosaic. As this mosaic grows by mitotic addition of new photoreceptors at the rim of the hemispheric retina, topological defects, called “Y-Junctions”, form to maintain approximately constant cell spacing. The generation of topological defects due to growth on a curved surface is a distinct feature of the cone mosaic not seen in other well-studied biological patterns like the R8 photoreceptor array in the *Drosophila* compound eye. Since defects can provide insight into cell-cell interactions responsible for pattern formation, we characterize the arrangement of cones in individual Y-Junction cores as well as the spatial distribution of Y-junctions across entire retinae. We find that for individual Y-junctions, the distribution of cones near the core corresponds closely to structures observed in physical crystals. In addition, Y-Junctions are organized into lines, called grain boundaries, from the retinal center to the periphery. In physical crystals, regardless of the initial distribution of defects, grain boundaries can form via the mobility of individual particles. By imaging in live fish, we demonstrate that grain boundaries in the cone mosaic instead appear during initial mosaic formation, without requiring defect motion. Motivated by this observation, we show that a computational model of repulsive cell-cell interactions generates a mosaic with grain boundaries. In contrast to paradigmatic models of fate specification in mostly motionless cell packings, this study emphasizes the role of cell motion, guided by cell-cell interactions during differentiation, in forming biological crystals. Such a route to the formation of regular patterns may be especially valuable in situations, like growth on a curved surface, where long-ranged, elastic, effective interactions between defects can help to group them into grain boundaries.

**AUTHOR SUMMARY:** From hair cells in the mammalian inner ear to the bristles on a fly’s back, sensory cells often form precise arrays, ensuring that these cells are evenly spread out on the tissue’s surface. Here we consider the zebrafish cone mosaic, a crystal of cone photoreceptors in the outer retinal layer. Because the cone mosaic grows from the rim of the curved retinal surface, new rows of cones (*i.e.*, defects) are inserted to maintain constant spacing between sensory cells. We study the spatial distribution of these defects to gain insight into how the cone pattern forms. By imaging retinae in live fish, we find that as differentiating cones are incorporated into the mosaic, defects form lines (grain boundaries) that separate mostly defect-free domains. Then, we show that a computational model based on repulsion between mobile cells during their incorporation into the mosaic generates similar grain boundaries. This study thus suggests that cell motion governed by repulsive cell-cell interactions can play an important role in establishing regular patterns in living systems.

## INTRODUCTION

In epithelial sheets that sense an external stimulus, the sensory function often depends on the spatial ordering of the constituent cells. In several examples [1–9], the pattern is sufficiently precise that if one knows the fate of just one cell, one can determine the identities of all the others. It remains a major challenge to understand how these extraordinarily regular cell arrays are created during development. Here we focus on one such system, the photoreceptor cell layer in the zebrafish retina, in which cone photoreceptors are organized by spectral subtype into a crystalline, two-dimensional lattice called the cone mosaic [10–12]; in particular, we use defects in this lattice as a window into possible mechanisms of mosaic formation. Although the precise evolutionary advantage and functional significance of the cone mosaic remains unknown, establishing an organized lattice in which each cone maintains some characteristic spacing from neighboring cones of the same subtype is thought to optimize sensitivity to a broad range of wavelengths over the full spatial extent of the retina [13–14].

Four spectral subtypes form the zebrafish cone mosaic: Red, Green, Blue, and Ultraviolet (UV) [15–16]. The ‘unit cell’, or the smallest repeating unit necessary to build the entire lattice, is composed of one Blue cone, one UV cone, two Green cones, and two Red cones (Fig. 1A-B). Blue and UV cones form interpenetrating anisotropic triangular sublattices (Fig. 1D). Green and Red cones form interpenetrating anisotropic honeycomb sublattices (Fig. 1E). Along ‘rows’, Blue cones alternate with UV cones, and Red cones alternate with Green cones (Fig. 1A-B). Along ‘columns’, each Blue cone is flanked by two Red cones, and each UV cone is flanked by two Green cones (Fig. 1A-B). Rows radiate from the center of the retina to the periphery. Columns are approximately parallel to the rim of the retina (Fig. 1A-C).

**Figure 1.**
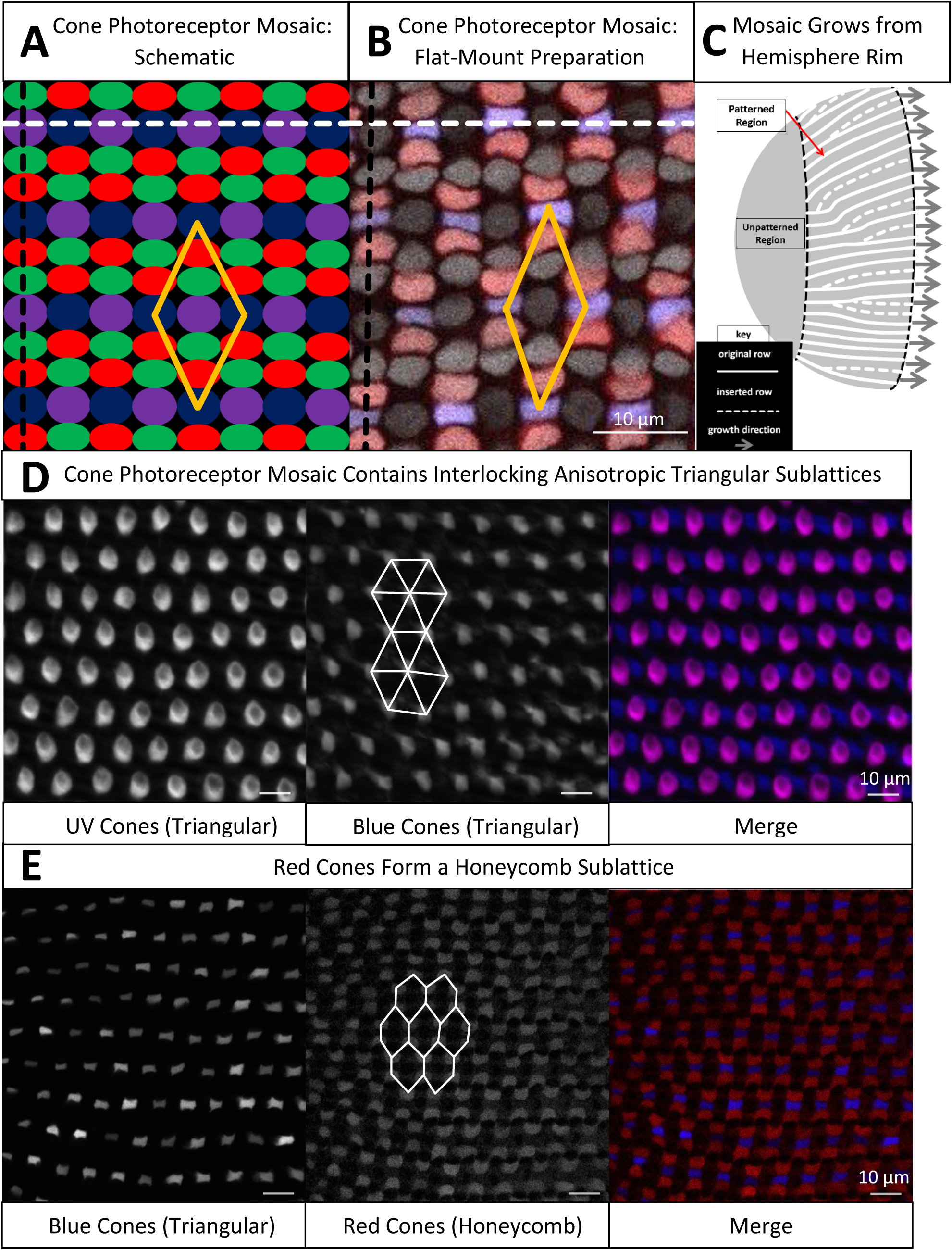
The cone mosaic is composed of four interpenetrating sublattices: two anisotropic triangular sublattices and two anisotropic honeycomb sublattices. **A)** Schematic of cone photoreceptors in the apical plane of the zebrafish retina. Cones are colored according to subtype: Red, Green, Blue, and UV (magenta). The ‘unit cell’ of the cone mosaic, which contains one UV cone, one Blue cone, two Green cones, and two Red cones, is indicated by the yellow parallelogram. The ‘row’ axis is indicated by the white dashed line, and the ‘column’ axis is indicated by the black dashed line. B) Cone mosaic from a flat-mount retinal preparation of an adult, triple transgenic fish, Tg[*sws2:GFP*; *trβ2:tdTomato*; *gnat2:CFP*]. Blue cones express a fluorescent reporter (pseudo-colored blue) under the control of *sws2* (Blue opsin) promoter, and Red cones express a fluorescent reporter (pseudo-colored red) under the control of the *trβ2* promoter. All cones express an additional fluorescent reporter (pseudo-colored gray) under the control of the *gnat2* promoter. Note that although UV and Green cones are not expressing different fluorescent reporters, we can distinguish between these two subtypes based on morphological differences. C) Schematic of photoreceptor epithelium, which lines the outer surface of the hemispheric retina. The central retina, which surrounds the hemispheric pole and forms during the larval period, is not patterned. As the retina grows by the mitotic addition of new photoreceptors (and other retinal cells) at the hemispheric rim (as indicated by gray arrows), there is a disorder-to-order transition (indicated by the black dashed line). After the disorder-to-order transition, the cone mosaic continues to grow by neurogenesis at the hemispheric rim throughout the life of the fish. Because the circumference of the hemisphere grows, rows of cells must be inserted to maintain approximately constant cell spacing. D) UV and Blue cones in a flat-mount retinal preparation from a double transgenic (Tg[*sws1:GFP*; *sws2:mCherry*]) line in which UV and Blue cones express distinct fluorescent reporters. UV cones (pseudo-colored magenta) form an anisotropic triangular sublattice that interpenetrates with an anisotropic triangular sublattice of Blue cones (pseudo-colored blue). We connect a subset of nearest neighbors in the Blue cone sublattice with solid white lines to aid in visualization of the pattern. E) Blue (pseudo-colored blue) and Red (pseudo-colored red) cones in the same flat-mount retinal preparation as panel B. Red cones neighbor Blue cones in each column. The Red cones form an anisotropic honeycomb sublattice. We connect a subset of nearest neighbors in the Red cone sublattice with solid white lines; note the different nearest neighbor patterns in the Blue cone triangular lattice (panel D) and the Red cone honeycomb lattice (panel E). The Green cones also form a honeycomb sublattice (not shown here).

The retinal hemisphere grows outward from the rim by mitotic addition of new photoreceptors (and other retinal cells) (Fig. 1C) [17–20]. Until approximately two to three weeks post-fertilization, the newly incorporated cones are not arranged in an ordered mosaic [21–22]. Then, a disorder-to-order transition, in which newly incorporated cones begin to form regular lattice, occurs. The region of cones generated earlier than this transition, called the ‘larval remnant’, remains disordered [21–22]. We call the rows that originate from the boundary of the larval remnant the ‘original rows’ (Fig. 1C).

As new cells are incorporated at the rim, the circumference of the retinal hemisphere enlarges, and the spacing between the original rows necessarily increases (Fig. 1C). To maintain approximately constant spacing between rows, new rows, that do not originate at the larval remnant, are inserted (Fig. 1C). The topological defects that generate new rows are called Y-Junctions [22–23]. For a crystal on a spherical (closed) surface, defects are inevitable, as required by Euler’s formula [24–27]. In contrast, defects in the hemispheric photoreceptor layer, a non-closed surface, result not from a fundamental topological constraint but from the biophysical requirement to maintain reasonable cell sizes and not to leave gaps between cells in the retinal epithelium.

The generation of topological defects to maintain approximately constant cell spacing during growth on a curved surface makes the cone mosaic distinct from other patterned tissues, such as sensory bristles [28] and R8 photoreceptors in *Drosophila* [1–7]. Previous investigators have noted the existence of defects in the teleost cone mosaic [22–23]. Because these topological defects can provide insight into the biological mechanisms of pattern formation [29–31], in this paper we characterize the spatial distribution of each cone subtype in the Y-Junction core and compare Y-junction cores to defect cores in physical crystals. We show that a Y-Junction is a dislocation [32–33], the insertion of a row and a column.

Additionally, we characterize the spatial distribution of Y-Junctions in the retinae. We demonstrate that the spatial distribution of Y-Junctions is as expected in a physical crystal near an energy minimum on a hemisphere [26, 34–39]. As in a physical crystal, the defects form lines, called grain boundaries, from the center of the retina to the periphery [26, 34–39]. In a physical crystal at finite temperature, defects are mobile; therefore, defects can coalesce into grain boundaries after formation of the crystal, regardless of the initial spatial distribution of defects [32-33, 37, 39-40]. We demonstrate that in the zebrafish retina, in contrast, grain boundaries appear during initial mosaic formation and do not require subsequent defect motion.

Having observed grain boundaries in fish retinae, we seek to take advantage of this finding to gain insight into the mechanisms of cone mosaic formation. We previously reported that cones of different subtypes are in approximately correct locations relative to each other within hours after they are generated by the terminal divisions of progenitor cells [41]. Though previous studies have documented interactions between cones in mature columns [21, 42], little is known about the mechanisms by which premature columns initially form; in particular, the genetic and signaling networks that lead to spectral fate specification remain almost completely unexplored. Evidence from embryonic retina suggests that the spectral subtype of each cone is determined at the time of a symmetric, terminal division of its precursor [43]. If this finding from embryonic retinae holds for juvenile and adult retinae, it implies that the two daughter cells of the same subtype must move away from each other after their birth in order to reach the correct positions in the cone mosaic. This would suggest that interactions between differentiating cones with an established subtype generate crystalline order as these cones are incorporated into the retina.

Inspired by this evidence from embryonic retinae as well as other examples of neural cell mosaics [44–48], we propose a computational model in which fate-committed cells repel each other in an anisotropic medium. This model generates grain boundaries during initial mosaic formation, consistent with our observations of fish retinae. We, then, contrast our model of motile, fate-committed cells with a second model in which cells are neither fate-committed nor motile. In this second model, inspired by the example of Notch-mediated lateral inhibition in neural fate specification, static cells in a disordered packing signal to each other at short range to set up a fate pattern [2–7, 28]. Because the signaling range is approximately equal to the cell size, we find that in the absence of cell motion, this mechanism of cell-cell signaling generates many excess defects (consistent with a very recent, independent study [9]). We conclude that our model of motile, fate-committed cells is more consistent with observations of cone mosaic formation than a model of cell-cell signaling in a disordered packing.

The biological example of grain boundary formation during initial patterning in zebrafish retinae also poses interesting physical questions. A primary concern in the existing physics literature has been the existence of grain boundaries in the ground state of crystals on curved surfaces [24-26, 34, 49-50]; although some aspects of the kinetics of crystal growth have also been considered [51–52], the question of how the growth geometry affects the positioning of defects has received little attention [40, 53]. For example, in which growth geometries does crystallization produce defect distributions that are close to the ground state without defect motion? We show that for crystal growth in geometries comparable to the zebrafish retina, repulsive cell-cell (more generally, particle-particle) interactions produce just such low energy defect distributions during the initial growth process.

In the remainder of this paper, after characterizing the spatial distribution of each cone subtype in the Y-Junction core, we demonstrate the presence of grain boundaries in fish retinae. To quantify whether grain boundary formation occurs via defect motion, we track motion of individual defects in the retina in live fish. By comparing the timescales of defect motion and grain boundary growth, we conclude that grain boundaries form as cones are initially incorporated into the mosaic. We explain why cone mosaic formation is unlikely to occur via fate specification in a static, disordered cell packing, and we test a model of cell motion guided by cell-cell repulsion in an anisotropic medium. The latter model generates grain boundaries during initial mosaic formation, consistent with our observations of the retina.

## RESULTS

### A Y-Junction is the insertion of a row and a column in the cone mosaic

To maintain approximately constant cell spacing as the retina grows by mitotic addition of cone photoreceptors at the rim, rows must be inserted (Fig. 1C) [22–23]. It is straightforward to demonstrate that a simple row insertion causes a disruption, that is not limited to a point defect but extends along an entire line, in the cone mosaic (Fig. 2A). To avoid this disruption along an entire line of the cone mosaic, it is necessary to consider more complex defects: the insertion of two rows (Fig. S1) or the insertion of a row and a column, neither of which disrupts formation of the cone mosaic. In the zebrafish cone mosaic, the most common topological defect is the insertion of a row and a column, *i.e.*, a ‘Y-Junction’ (Fig. 2B-C). To understand why the insertion of a row and a column is expected to be the most prevalent defect, we employ a tool used in analyzing defects in physical crystals: the Burgers vector [32–33].

**Figure 2.**
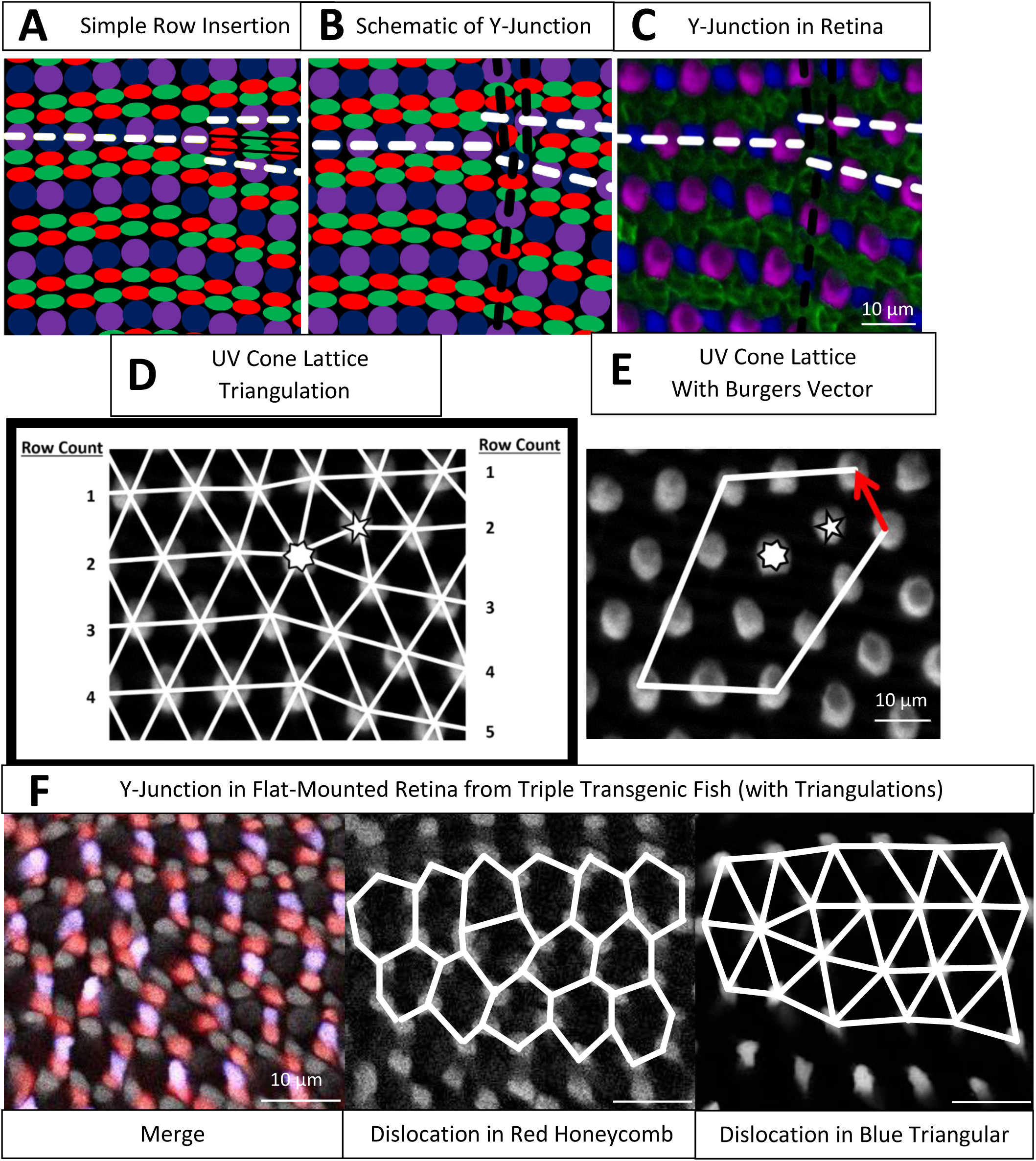
A Y-Junction, a topological defect in the cone mosaic, is an insertion of a row and a column. **A)** Schematic of simple row insertion in cone mosaic. The rows associated with the defect are indicated by the white dashed lines. As new cone photoreceptors are incorporated to the right of the defect, a series of improper cone contacts, indicated by the black box, within columns form. If the colors were reversed in one of the new columns, so that a Red cone contacts a Green cone, there would instead be improper Red-UV and Blue-Green contacts. **B)** Schematic of a Y-Junction, a topological defect in the zebrafish cone mosaic. The rows associated with the defect are indicated by white dashed lines, and the columns associated with the defect are indicated by black dashed lines. A Y-Junction, the insertion of a row and a column, only disrupts the cone mosaic near the core rather than along an entire line of contacts. **C)** A Y-Junction in a flat-mount retinal preparation from an adult, double transgenic (Tg[*sws1:GFP*; *sws2:mCherry*]) line in which UV and Blue cones express distinct fluorescent reporters (pseudo-colored magenta and blue, respectively) under the control of the UV and Blue opsin promoters, respectively. Red and Green cones (both pseudo-colored green) are visualized by antibody staining. **D)** Each UV cone from panel C is connected to its nearest UV cone neighbors with white bonds. The seven-coordinated UV cone is indicated by a seven-sided star, and the five-coordinated UV cone is indicated by a five-sided star. To the left and right of the defect, the rows are counted. **E)** A circuit of triangulation bonds is drawn around the defect from panels C-D. The red arrow corresponds to the Burgers vector, the additional bond that is necessary to close the circuit when the circuit contains a dislocation. **F)** Y-Junction in the same flat-mount retinal preparation as in Fig. 1B. The cells with a round morphology and dim fluorescence are UV cones. The Red cones are pseudo-colored red. The Blue cones are pseudo-colored blue. The remaining cones (bright gray fluorescence) are Green cones. We connect nearest neighbors in the Red cone sublattice and Blue cone sublattice with solid white lines.

#### Y-Junction generates minimal lattice deformation, as quantified by Burgers vector

As discussed in the introduction, the unit cell of the cone mosaic is composed of one Blue cone, one UV cone, two Green cones, and two Red cones (Fig. 1A-B). One can generate an infinite cone mosaic on a flat plane given the unit cell and two lattice vectors, which define the Bravais lattice [54]. The Bravais lattice defines which defects one expects to observe in the cone mosaic lattice, though the distribution of particles in the defect core may vary [32–33, 54]. For the sake of clarity, we analyze the defects in the cone mosaic from the perspective of a cone subtype that appears only once in the unit cell: UV cones.

To define the Burgers vector, we build a triangulation for the UV cones in which we connect nearest neighbors of the same cone subtype (Fig. 2D-E). Away from the core of the defect, every UV cone is surrounded by six nearest UV cone neighbors [32–33, 54]. Near the defect core, as in physical crystals that are triangular, one UV cone is surrounded by seven nearest UV cone neighbors. Neighboring this seven-coordinated UV cone is a UV cone which has only five nearest UV cone neighbors. This pair of five- and seven-coordinated UV cones constitutes the core of the dislocation.

We, then, construct a circuit that surrounds the core of the defect. If there were no defect, the circuit would be a parallelogram (Fig. 2E). The bottom side of the circuit would contain as many bonds as the top side of the circuit. The right side of the circuit would contain as many bonds as the left side of the circuit. If there is a dislocation inside of the circuit, to close the circuit, one must add a bond, called the Burgers vector [32–33]. The magnitude of the Burgers vector quantifies the amount of lattice deformation associated with the dislocation [32–33].

In physical crystals, where the elastic deformation associated with the dislocation is proportional to the magnitude of the Burgers vector squared, the defect that generates the least deformation is expected to be the most prevalent. Even though we have no reason *a priori* to treat this biological crystal as elastic, we expect this measure of deformation to be generally applicable. The mechanism that drives ordering in a non-physical crystal likely also resists large deformations due to defects in the lattice.

The Y-Junction in the cone mosaic lattice is the dislocation that introduces the smallest deformation. This can be seen by comparing the Burgers vector of a Y-Junction to the Burgers vector of a double row insertion. For a double row insertion, the length of the Burgers vector is equal to the spacing between UV cones along a column, which is approximately twelve and a quarter microns, as compared to the Burgers vector of a Y-Junction, with a length of approximately eight microns (for quantification of spacings between UV cones in same column and in same row, see Fig. S2). For the sake of minimizing lattice deformations, we expect double row insertions to be less prevalent than Y-Junctions.

#### Distribution of Red and Green cones near the Y-Junction core

For Red and Green cones, that each appear twice in the unit cell, we connect nearest neighbors of the same subtype and analyze the spatial distribution in the Y-Junction core. Away from a Y-Junction core, the cells that form the lattice can be grouped into hexagons (Figs. 1E, 2F, S3) [54–56], but this grouping breaks down near the defect core. The distribution of Red and Green cones in the Y-Junction core is variable but is often either a ‘glide’ dislocation or a ‘shuffle’ dislocation, which differ in the distributions of cones at the core.

**Figure 3.**
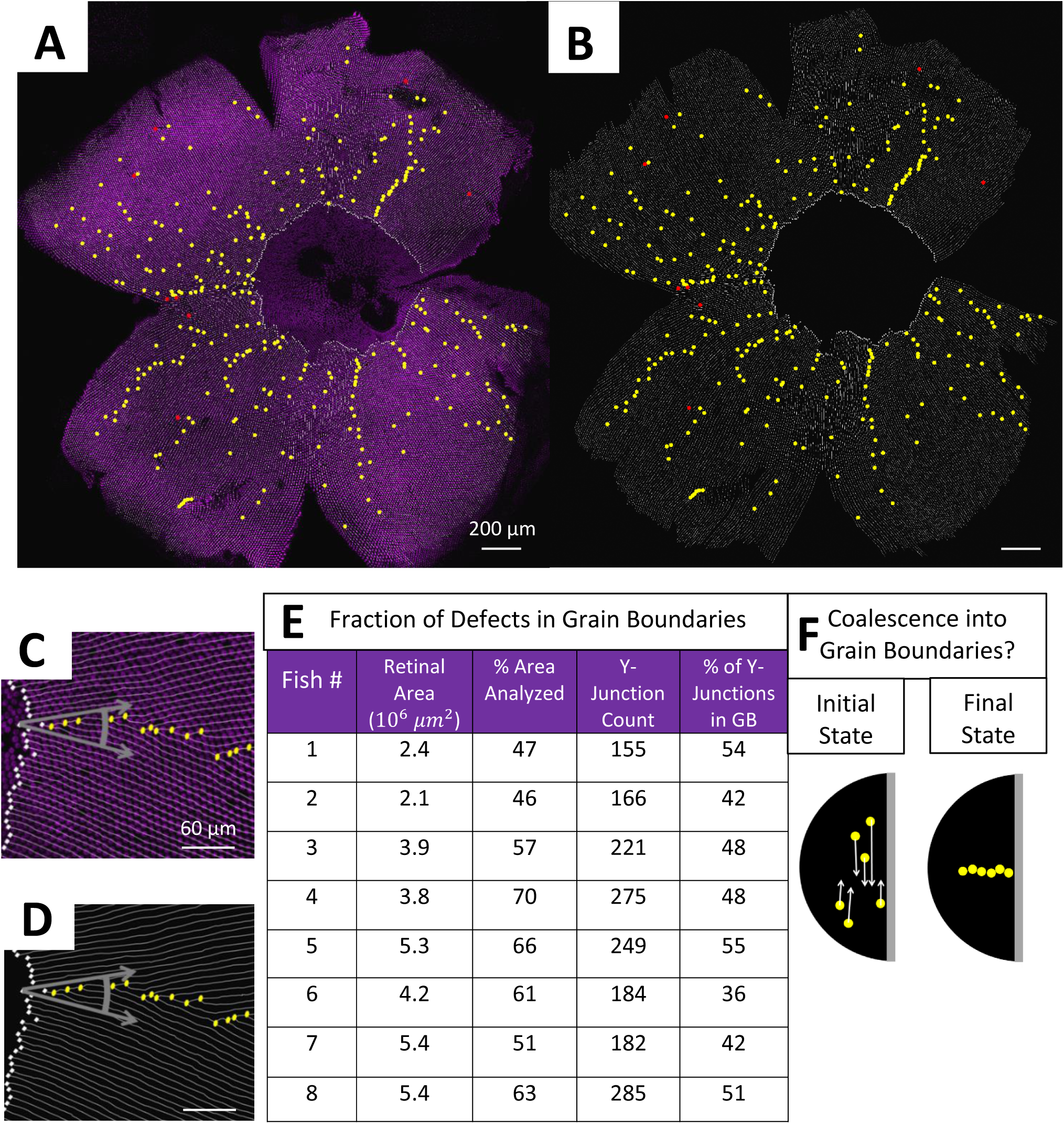
Y-Junctions form lines, called grain boundaries, that run from the center of the retina to the periphery. **A)** Flat-mounted retina in which UV cones express a transgenic reporter (pseudo-colored magenta) under control of UV cone opsin promoter. Rows of UV cones (white lines) are traced. The yellow dots correspond to Y-Junction. The red dots correspond to reverse Y-Junction, generating row deletions. White squares indicate the onset of pattern where we began tracing rows. The dorsal side of the retina is left, and the ventral side is right. The temporal side is down, and the nasal side is up. **B)** Row tracing and identification of defects from the retina in panel A. **C)** Example of a grain boundary from the retina in panel A. The gray arrows indicate that a grain boundary reconciles domains of differing crystallographic orientation. **D)** Grain boundary presented in panel C with only the row tracing. **E)** From the row-tracing of seven retinae, we show the percentage of retinal area analyzed and the number of Y-Junctions identified. We show the fraction of defects in grain boundaries (see Methods). **F)** Illustration of potential role of defect motion in generating final spatial configuration of defects. The black region is the photoreceptor epithelium, and the gray region indicates the margin from which the photoreceptor epithelium grows. The yellow circles denote Y-Junctions. If defect motion does occur in the cone mosaic, it could allow defects to line up into grain boundaries. If defect motion is too slow, the patterning mechanism would have to generate grain boundaries during initial mosaic formation.

A ‘glide’ dislocation has a heptagon and a pentagon in the core. For example, in Fig. S3A-B, one heptagon of Red cones neighbors a pentagon of Red cones. A ‘shuffle’ dislocation has a single octagon in the core. For Red cones (Fig. S3E), the octagon contains two UV cones, and for Green cones (Fig. S3D, F), the octagon contains two Blue cones. Interestingly, both ‘glide’ and ‘shuffle’ dislocations are commonly observed in honeycomb crystals like graphene [55–56].

### Y-Junctions form lines, called grain boundaries, that run from center of retina to periphery

Having verified that individual Y-Junctions are the dislocations that generate minimal lattice deformation, we next study the spatial distribution of Y-Junctions on the retinal surface. On the retinal hemisphere, the row direction rotates by 2π about the pole of the hemisphere, similar to the convergence of longitudinal lines toward a pole on the globe (Fig. 1C). For physical crystals in this orientation, the ground state contains lines of dislocations, called grain boundaries, from the center (pole) of the hemisphere to the edge (equator) [26, 57]. In physical crystals, dislocations are mobile; therefore, in physical crystals, it is possible for defects to rearrange into grain boundaries after crystallization, regardless of the initial spatial distribution of defects [32-33, 37, 39-40, 57].

In a biological crystal, it is not obvious that the Y-Junctions will form a spatial pattern that is equivalent to the ground state of a physical crystal. If Y-Junctions do form grain boundaries, however, we may be able to leverage that information to understand the mechanism by which the biological crystal forms. By manually tracing rows of UV cones over approximately fifty percent of the retinal area in eight retinae (see Methods), we identified the locations of approximately one thousand seven hundred Y-Junctions.

#### In flat-mounted retinae, a large fraction of Y-Junctions form grain boundaries

Y-Junctions do, indeed, form grain boundaries that run from the center of the retina to the periphery (Figs. 3A-B, S4). These grain boundaries reconcile domains of differing crystallographic orientations (Fig. 3C-D). The angle by which the local row direction rotates from one side of a grain boundary to the other side is determined by the linear density of Y-Junctions in the grain boundary and the length of the Burgers vector of an individual Y-Junction [32–33, 57].

**Figure 4.**
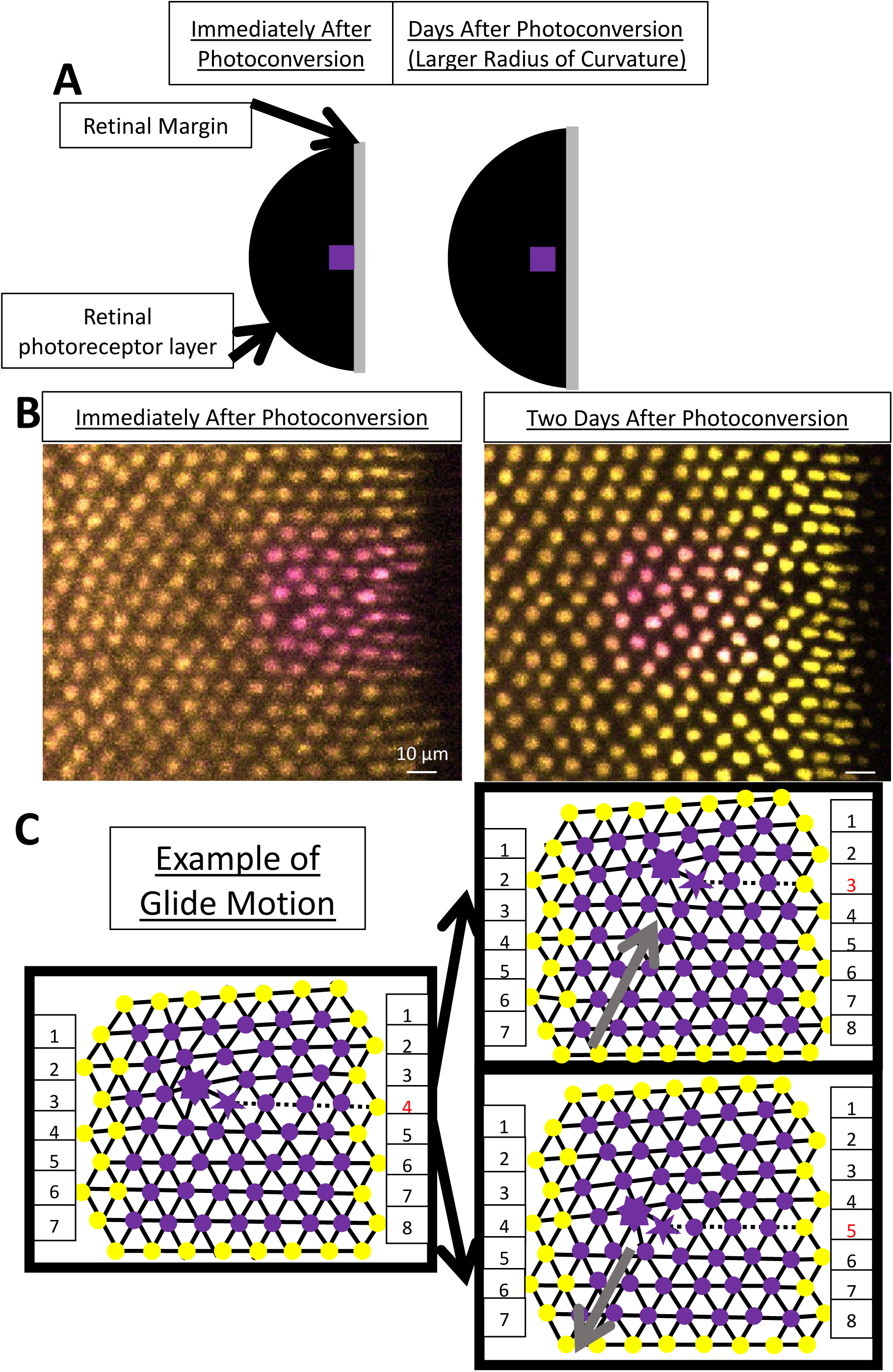
By photoconverting UV cones near the retinal margin, we track Y-Junction motion. **A)** Schematic of photoconverted UV cones in the photoreceptor epithelium near the retinal margin. We photoconvert a patch of UV cones (indicated by the purple box) near the retinal margin, where new UV cone photoreceptors are incorporated by mitotic addition. After a certain amount of time (two, three or four days) during which the retina has grown, we image the photoconverted region. **B)** Example of patch of UV cones immediately after photoconversion and two days later. In this line (Tg[*sws1:nEOS*]), UV cones express a nuclear-localized, photoconvertible fluorescent protein under control of the UV cone opsin promoter. The non-photoconverted fluorescent protein is pseudo-colored yellow, and the photoconverted fluorescent protein is pseudo-colored magenta. The retinal margin is to the right of each image. Note that approximately eight columns of UV cones have been added in the two days since photoconversion. **C)** Glide motion involves subtle motion of individual UV cones near the Y-junction core. The magenta circles correspond to UV cones with photoconverted fluorescent signal, and the yellow circles correspond to the surrounding UV cones with non-photoconverted fluorescent signal. Every cone is connected to nearest neighbors in the triangulation. The five-sided and seven-sided star indicate the five-coordinated and seven-coordinated UV cones, respectively. The dashed black line corresponds to the “inserted” row. The two triangulations on the right describe the position of UV cones (from the left triangulation) after glide in the direction denoted by the gray arrow. Note that the assignment of the five- and seven-coordinated UV cones has shifted by one unit.

Although these grain boundaries are identifiable by eye, we developed an objective definition of grain boundaries, which can be applied to biological data and, later, to simulation results of potential models of cone mosaic formation. To count which defects are in grain boundaries, the measure tests whether a defect belongs to an approximately linear chain of at least four other defects, which are each other’s nearest neighbors (see Methods). By applying this measure to the eight analyzed retinae, we found that approximately fifty percent of the identified Y-Junctions are in grain boundaries (Figs. 3E, S4).

### Defect motion is not responsible for grain boundary formation

To study whether defect motion is responsible for the existence of grain boundaries, we need to observe the dynamics of individual cones during cone mosaic formation in live fish rather than the fixed positions of cones in flat-mounted retinae. If defect motion is not responsible for the existence of grain boundaries, we will be able to test potential models of cone mosaic formation based on their ability to form grain boundaries.

To quantify the motion of individual UV cones during cone mosaic development, one must observe the same region of the retina in the same fish at two distinct time points as the fish remains alive. To make sure that one can locate the same set of UV cones at the two distinct time points, one must have some common point or common boundary as a reference for UV cone positions in the two images.

To locate the same set of UV cones at two different timepoints in live fish, we use transgenic zebrafish in which the UV cones express a nuclear-localized, photoconvertible fluorescent protein under the control of the UV cone opsin promoter (see Methods). We photoconvert and image a small patch at the retinal margin, near where newly generated UV cones are incorporated into the growing retina (Fig. 4A-B). At the time of imaging, the cone mosaic is composed of approximately eighty columns and is growing by approximately three columns of cones per day [41]. Two, three, or four days later, we image both photoconverted and non-photoconverted UV cones in the same retinal area (Fig. 4A-B).

#### Eliminating the possibility of defect motion perpendicular to the Burgers vector

To understand what types of motion we expect to observe, it is useful to revisit the concept of a Burgers vector. When a dislocation moves along the direction defined by the Burgers vector, this motion requires only subtle rearrangements of individual particles, here UV cones, near the core of the dislocation. This motion is called glide motion [32–33]. To illustrate this motion in Fig. 4C, we denote the inserted row associated with the Y-Junction as well as the five- and seven-coordinated UV cones. When the dislocation glides by one unit, as illustrated in Fig. 4C, the initial inserted row incorporates itself into a neighboring row, as a new inserted row is generated. The assignment of the five- and seven-coordinated UV cones shifts by one unit in the direction of the glide motion. Glide by one unit is the flipping of one bond in the UV cone triangulation near the Y-Junction core.

On the other hand, motion of dislocations perpendicular to the direction defined by the Burgers vector, called glide motion, requires the creation or annihilation of point defects, which are interstitials or vacancies in the crystal [32–33]. A vacancy in the cone mosaic corresponds to the absence of six cones in the same site, which we never observe (Fig. S5). Therefore, in monitoring the motion of individual UV cones, we test specifically for glide motion rather than climb motion [37].

#### Quantifying glide motion

Based on the positions of UV cones, we connect nearest neighbors in the lattice (triangulation method described in Methods). We identify the location of the Y-Junction core based on the location of the inserted row (Fig. 5A). To quantify glide motion, we search for bond flips (Figs. 4C, 5A) along the glide line (see Methods) near the Y-Junction core between the time of photoconversion and the time of subsequent imaging (two, three, or four days later).

**Figure 5.**
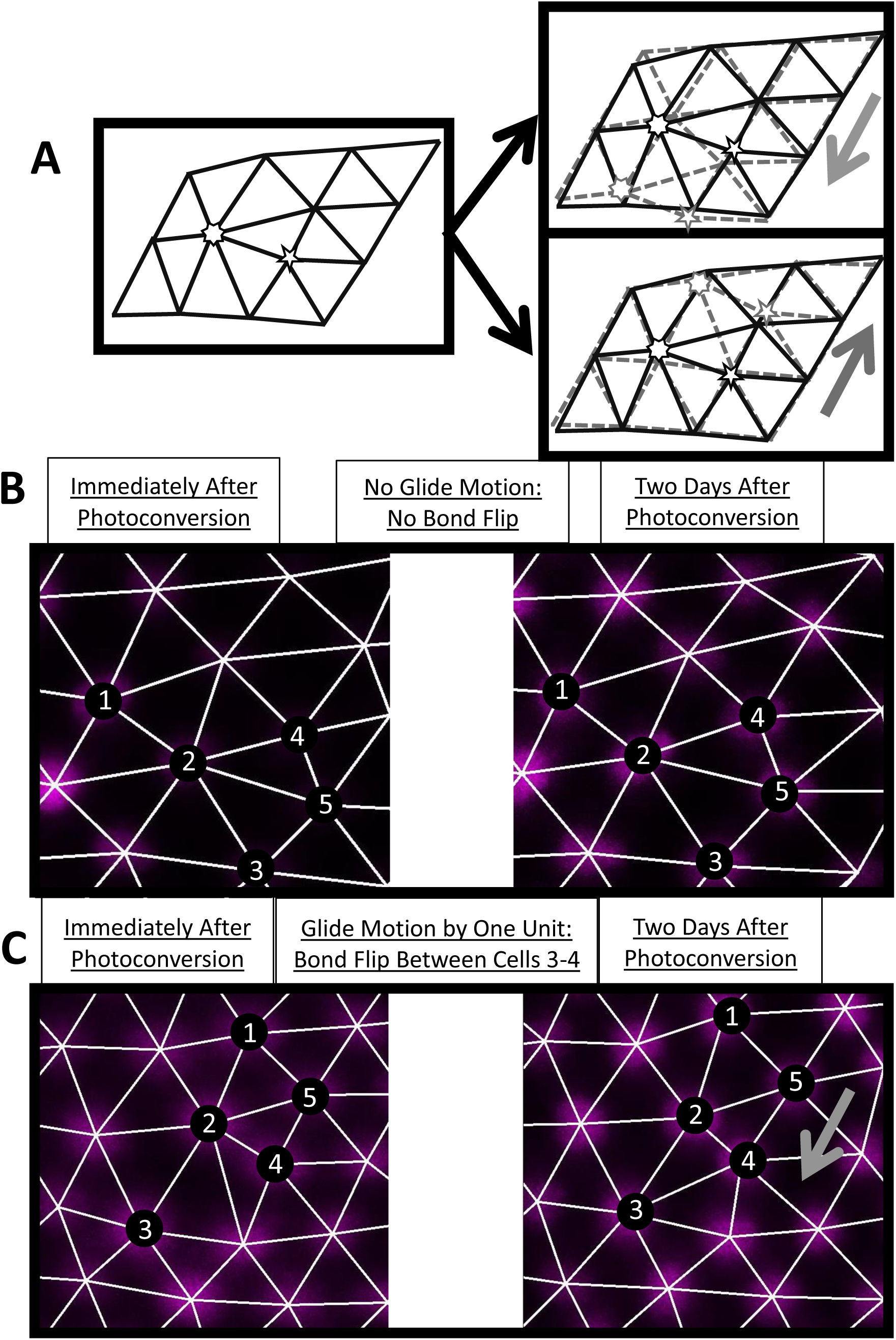
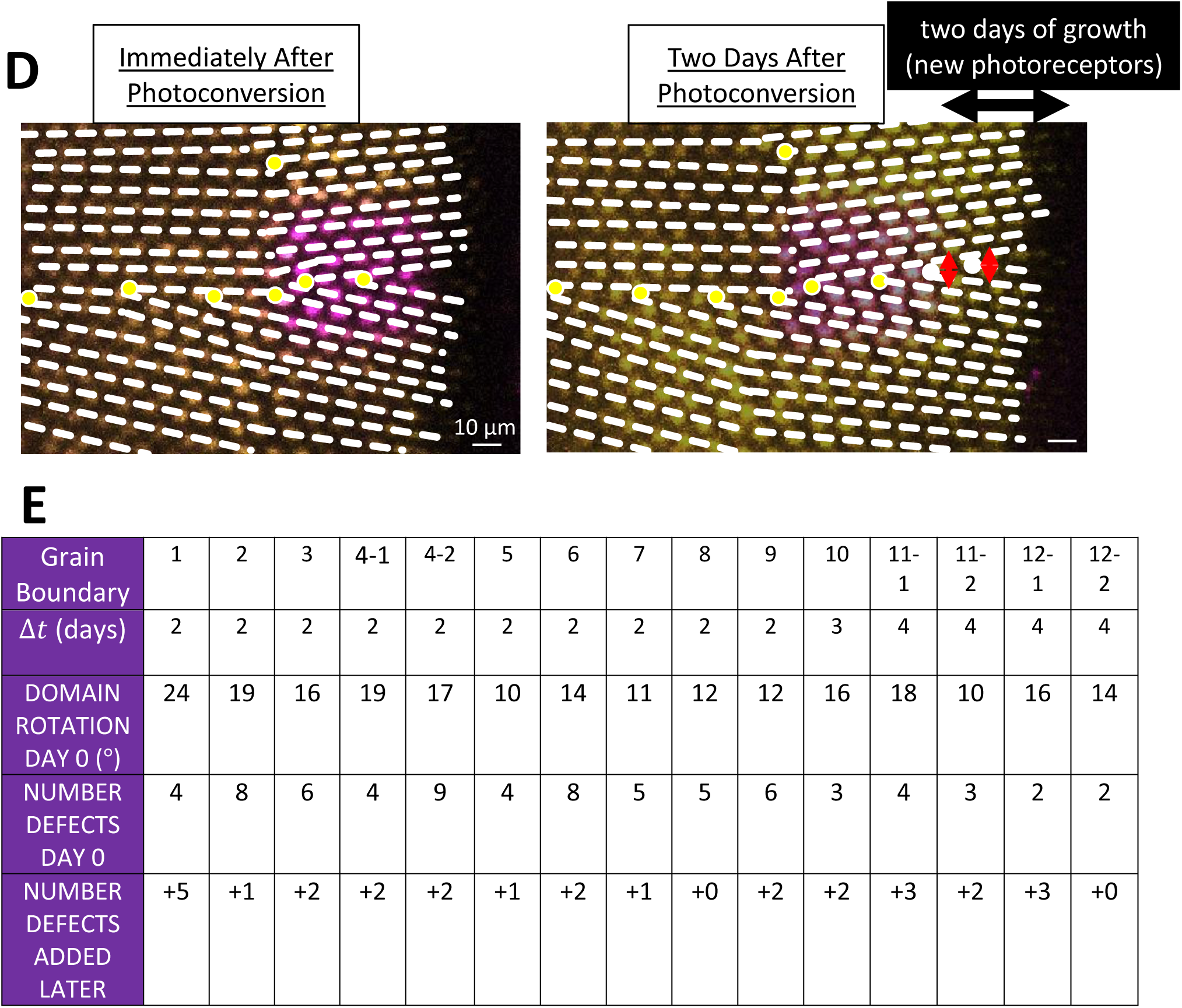
By estimating the timescale of Y-Junction motion, we conclude that Y-Junctions line up into grain boundaries during initial mosaic formation rather than by subsequent Y-Junction motion. **A)** Expected motion of individual UV cones in the case of glide motion by one unit in either direction. The triangulation on the left corresponds to UV cones near the core of a defect. There is a UV cone on each site of the triangulation. The two triangulations on the right describe the positions of UV cones (from the left triangulation) after glide in the direction denoted by the gray arrow. The UV cones (originally sitting on the sites of the black triangulation) shift to the sites of the gray triangulation. Note that the originally five- and seven-coordinated UV cones in the black triangulation both become six-coordinated. **B)** Example of Y-Junction within photoconverted region in which no bond flips over the course of two days. The photoconverted fluorescent signal in UV cone nuclei is pseudo-colored magenta. The white lines are the triangulation of the UV cones. For reference, the same exact five cones are labeled in both images. **C)** Example of Y-Junction within photoconverted region from Fig. 4B. One bond has flipped in the triangulation over two days, meaning that the Y-Junction has glided in the direction of the gray arrow by a unit. **D)** Observation of the growth of a grain boundary during initial mosaic formation. Immediately after photoconversion, one observes seven Y-Junctions (indicated by yellow dots), six within a grain boundary and an isolated Y-Junction nearby. Rows of UV cones are traced with white dashed lines. Two days later, one observes two additional Y-Junctions incorporated into the grain boundary. Based on the constraint that a Y-Junction does not glide faster than one unit in two days, the Y-Junctions must have formed within the regions indicated by the red arrows (*i.e.*, had to form within the grain boundary). The black arrow indicates the columns of cones incorporated since the time of photoconversion. **E)** Growth of fifteen grain boundaries in live fish. In the image immediately after photoconversion, we measure how much the row direction rotates at the retinal margin (see Methods). For all fifteen cases in which the row direction rotates by more than ten degrees at the retinal margin, we count the number of defects in the corresponding grain boundary at the time of photoconversion. We, then, count the number of defects added to the grain boundary by the time of later imaging. Though we only photoconverted one region per fish, that region sometimes neighbors two grain boundaries, allowing us to measure growth of two grain boundaries in the same fish (*e.g.*, 4-1 and 4-2). The image in panel D corresponds to grain boundary 3.

We observe non-negligible motion near the core, but we never observe glide motion by more than one unit per two days, where glide motion by one unit is illustrated in Fig. 4C. We show two examples of Y-Junctions within photoconverted regions: an example in which there is no glide motion (Fig. 5B) and an example in which the Y-Junction glides by one unit in two days (Fig. 5C). These experiments provide an upper bound on the rate of glide motion (*i.e.*, one unit per two days). If we compare this constraint on the timescale of glide motion to the timescale of grain boundary formation, we can determine whether the presence of grain boundaries can plausibly be attributed to the coalescence of initially isolated dislocations that move together after the crystal forms.

#### Comparing the timescale of glide motion to the timescale of grain boundary formation

In many of our photoconversion experiments, we photoconvert patches of UV cones near at least one existing grain boundary (Figs. 5D, S6). At the time of subsequent imaging of the UV cones near the photoconverted region (*i.e.*, two to four days later), approximately eight UV cone columns are newly incorporated into the cone mosaic. After identifying Y-Junctions in the newly incorporated columns of UV cones, we ask whether their locations are correlated with the positions of existing grain boundaries (*i.e.*, observed at the time of photoconversion). If we do observe a correlation between the positions of new Y-Junctions and existing lines of Y-Junctions, we ask whether that correlation could be explained by Y-Junction motion.

**Figure 6.**
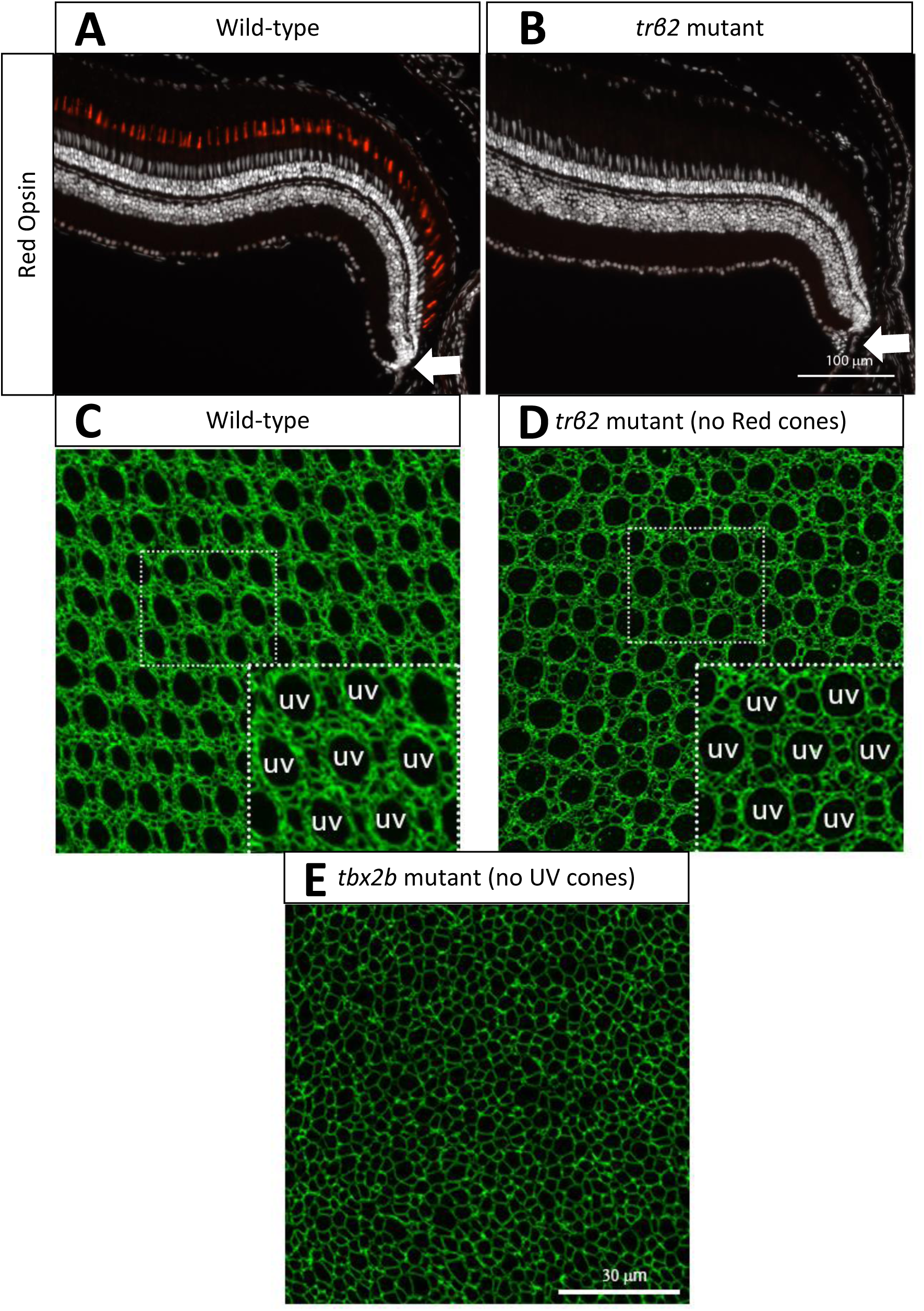
Cone mosaic formation is not disrupted in Red cone mutant but is disrupted in UV cone mutant. **A)** Cross-section of wild-type retina in which Red cones are labeled by immunostaining of Red cone opsin. White arrow indicates approximate location of precolumn area [41]. **B)** Cross-section of *trβ2* mutant in which Red cones are labeled by immunostaining of Red cone opsin. Please note the absence of differentiated Red cones. White arrow indicates approximate location of precolumn area. **C)** Apical plane of wild-type cone mosaic lattice in retinal flat-mounts in which cell profiles are stained by anti-ZO-1. UV cones are indicated in the inset. **D)** Apical plane of cone mosaic lattice in retinal flat-mount from *trβ2* mutant, which lacks Red cones. Cell profiles are stained by anti-ZO-1. UV cones (indicated in inset) are identified based on large, rounded profiles. Please note that the triangular lattice of UV cones is not disrupted in the absence of Red cones. **E)** Apical plane of cones in retinal flat-mount from *tbx2b* mutant, which lacks UV cones. The cone mosaic is disrupted in this mutant.

Out of the eighteen samples, twelve samples have grain boundaries near the retinal margin at the time of photoconversion. Since some samples have two grain boundaries, in total we observe fifteen grain boundaries in live fish (Figs. 5E, S6; see Methods). Two to four days later, we identify the positions of newly incorporated Y-Junctions. To these samples, we apply the following null model: each new Y-Junction’s initial position is uncorrelated with existing grain boundaries, but after formation, each new defect moves approximately one row unit closer to the closest grain boundary (see Methods). We find that newly incorporated defects are more aligned with existing grain boundaries than can be explained by this null model (p < 0.0001; see Methods). Because glide motion is too slow relative to the grain boundary growth, we conclude that grain boundaries form at the time of cone mosaic formation, not by subsequent defect motion.

### Testing Computational Models of Cone Mosaic Formation

Near the rim of the retinal hemisphere (Fig. 6A) is a region defined as the precolumn area, where newly generated but not fully differentiated cones are in approximately correct locations relative to each other based on cone subtype before the formation of mature columns [41]. Composed of differentiating post-mitotic cones, this precolumn area lies between two regions: a central region of mature columns and the rim, which contains proliferative cells. It remains unclear how proper positioning of cones by subtype in the precolumn area occurs, but importantly, this must occur within hours of the generation of post-mitotic cells by terminal divisions of neighboring proliferative cells [41].

In this section, we ask whether our observations of grain boundary creation can provide information about the mechanism for proper positioning of cones by subtype into immature columns. In principle, one can imagine two extremal models for the creation of regular cell fate patterns in biological tissues. In the first model, cell fates are specified first, and motile cells with a clear identity then rearrange themselves into the final pattern; in the second model, cells instead first arrange themselves in space, then the correct fate pattern is imposed on this static cell packing by cell-cell signaling. Evidence from embryonic retinae suggests that post-mitotic cones are of fixed subtype and move relative to each other during integration into the retina [43].

This finding from embryonic retinae, together with other examples of cell-cell repulsion in neural cell mosaics [44–48], suggests that mosaic formation in zebrafish retina falls closer to the first model. To test whether such a picture is consistent with the observed behavior of grain boundaries, we construct a computational model in which fate-specified cones repel each other during differentiation. After finding that this cell-cell repulsion model does indeed generate grain boundaries during the initial process of differentiation, we demonstrate that the alternative model, in which cell fate patterns arise through lateral inhibition in a static and disordered packing, is not likely to be responsible for cone mosaic formation. Before turning to the descriptions of the two models, we first gather some additional experimental data, on which cone subtypes are essential to establishing a crystalline mosaic and on lattice anisotropy in the zebrafish mosaic, to help in model formulation.

#### Absence of Red cones does not disrupt cone mosaic formation, but the absence of UV cones does

As noted above, crystals are described by identifying the smallest repeating unit that can be used to build the crystal, the unit cell – consisting of one Blue, one UV, two Red, and two Green cones (Fig. 1A) for the zebrafish retina – and the way different unit cells are positioned relative to each other in space, the Bravais lattice – for the zebrafish retina, a slightly anisotropic triangular lattice [32–33, 54]. It is well-established that the defect core structures can depend on all the features of the unit cell but that elastic interactions between two defects are determined only by the Bravais lattice and the defects’ Burgers vectors [25-26, 34, 54, 58]. Thus, we expect that most features of the spatial distribution of Y-junctions, which should depend primarily on defect-defect interactions [25-26, 34, 38, 49-50], can be recapitulated by a model in which each unit cell is replaced by a single cone photoreceptor. In order to provide biological justification for this simplification of the cone mosaic and to determine which cone subtype to focus on, we consider mutants in which certain cone subtypes are absent.

To evaluate the role of Red cones in establishing a crystalline mosaic, we generated a targeted mutation in a single gene that resulted in a loss of Red cones (Fig. 6B; see Methods). This gene, *trß2*, is an early fate marker of Red cones and is expressed in proliferating progenitors and mature Red cones, but not other cone subtypes [41, 43]. All other cone subtypes are still present in the outer retinal layer (Fig. S7A). Strikingly, in the *trß2* mutant, we find that cone mosaic formation, including ordering of UV cones, is not disrupted by the absence of Red cones (Fig. 6D).

The robustness of cone mosaic formation to the absence of Red cones is in marked contrast with previous experiments with *tbx2b* mutants in which UV cones are largely absent [59]. In *tbx2b* mutants, it is difficult to discern long-ranged crystalline order in the cone positions (Fig. 6E). The spatial distribution of cones and absence of long-ranged order is similar in other zebrafish mutants in which cell-cell adhesion is perturbed [21, 42]. Given this evidence from both *trß2* mutants and *tbx2b* mutants, we simplify the cone mosaic to a lattice of UV cones.

#### Measuring lattice anisotropy of the cone mosaic in live fish

Previous studies have established the importance of anisotropy of the mosaic lattice both in its formation and refinement [21, 41–43]. In modeling the cone mosaic lattice as a lattice of UV cones, we need to make sure that we produce a lattice with the same anisotropic spacing as the cone mosaic. For this reason, we first measure the lattice vectors of the UV cone sublattice in live fish. We use images of photoconverted regions, immediately after photoconversion, in which there are no Y-Junctions (Fig. S2).

In an isotropic triangular lattice, the ratio of the distance between UV cones in the column direction to the distance between UV cones in the row direction would be equal to the square root of three. We find that this ratio is approximately six-fifths in our live images of the UV cone sublattice (Fig. S2). This means that the spacing along the row direction is longer than would be expected in an isotropic triangular lattice. We use this degree of anisotropy as an input for modelling the cone mosaic formation.

#### Model of repulsive interactions between fate-committed cells generates grain boundaries during initial cone mosaic formation

In building a model of cone mosaic formation, we hypothesize that cell motion is generated by repulsive interactions between fate-committed cones of the same subtype as they differentiate in a mechanically anisotropic medium. This model is motivated by cell mosaics in other vertebrate retinal layers [44–47] and in the nervous system in *Drosophila* larvae [48]. For example, in mice, specific neuron subtypes disperse after fate commitment and during morphological differentiation [46–47]. By cell ablation, previous investigators have established that for retinal horizontal cells in mice, the cells are fate-committed and homotypically interact via transient neurites (*i.e.*, cell processes) [47]. Similarly, in *Drosophila*, certain classes of peripheral sensory neurons tile the body wall. By ablation of dendritic processes, previous investigators have established that neurons of a specific class (class IV) interact homotypically to establish a tiling of the body wall by non-overlapping cell territories [48].

To model repulsive interactions between fate-committed cones of the same subtype, leading to a preferred spacing in the row and column directions, we employ a phase-field model of crystallization (Fig. 7A-B). These models are widely employed to describe various processes in physical crystals, including nucleation of crystalline domains in a supercooled fluid, epitaxial film growth from a neighboring liquid phase, and the mechanical hardness of a solid based on the microstructure of crystalline domains [60–61].

Phase-field crystal models employ a continuum field, which corresponds to modulations in the particle density (Fig. 7B) [60–68]. Based on our simplification of the cone mosaic to a lattice of UV cones, the particle density represents the density of the UV cone subtype. If the field is homogeneous in space, the UV cones form a “liquid,” with no clear periodicity. If the field is periodic in space, the UV cones form an ordered crystal. The field starts out entirely uniform except near the center of the simulation domain (see Methods). The crystalline region grows outward from the center, as fate-committed cones in the disordered region are incorporated (Fig. 7A-B, Vids. 1-2).

**Figure 7.**
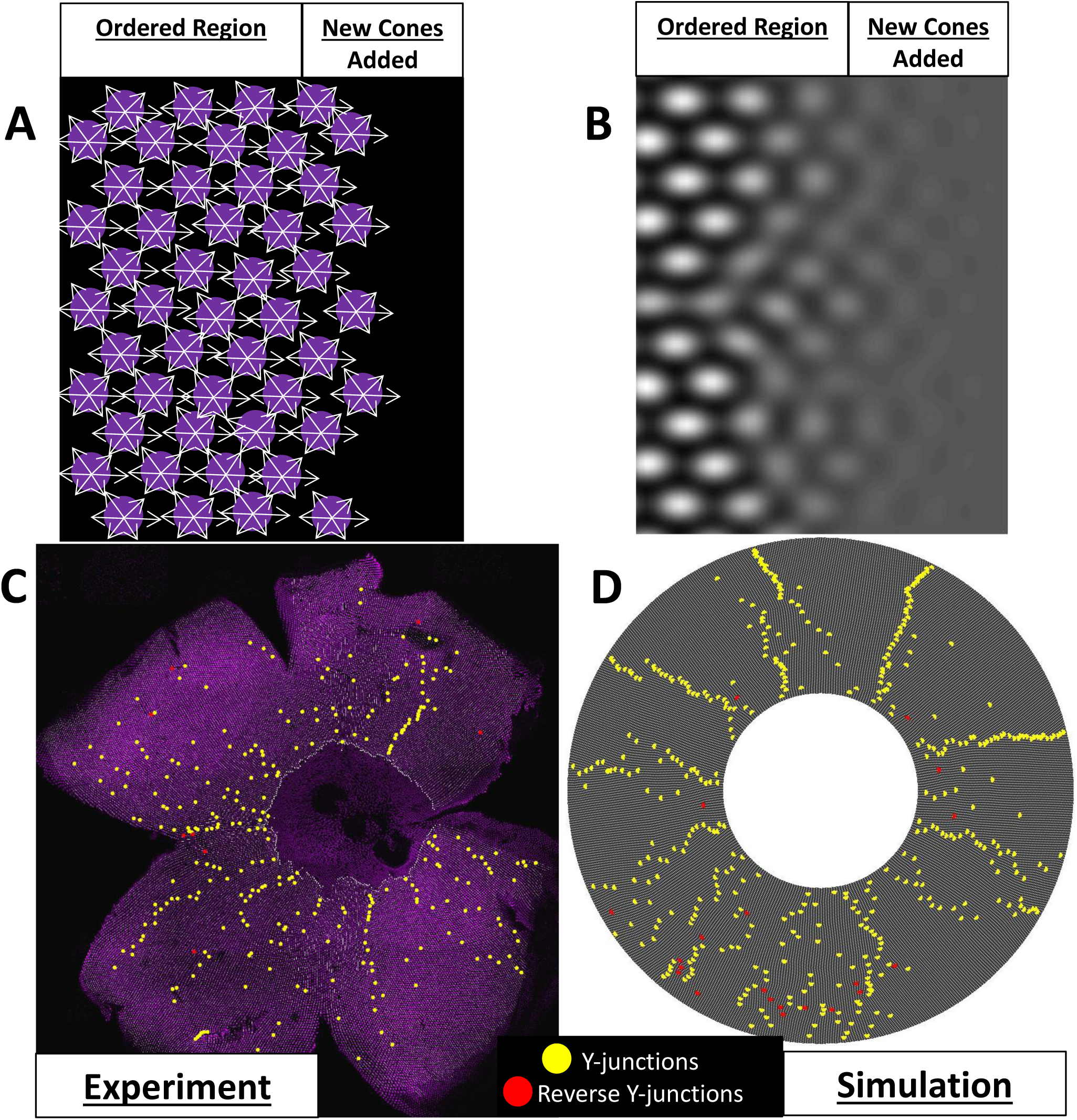
Phase-field crystal model of cone mosaic formation. **A)** Schematic of proposed contact-interaction model in which fate-committed cones interact homotypically, forming an anisotropic lattice. In this illustration, UV cones, denoted by magenta circles, interact with nearest neighbors of the same subtype. The contact interactions are denoted by the white arrows. New cones, which are already fate-committed, are incorporated to the right of the ordered region. **B)** In the phase-field crystal model, a continuum field describes the positions of, in this case, UV cones. Peaks in the density, here denoted by white regions, are regions in which one is likely to find a UV cone; troughs in the density, here denoted by dark regions, are regions in which one is unlikely to find a UV cone. Gray regions correspond to regions in which there is a lack of ordering in positions of UV cones. In this case, those gray regions correspond to the area in which new terminally differentiated UV cones will be incorporated into the existing crystal. **C)** Image of a flat-mounted retina in which UV cones express a transgenic reporter under the control of the UV cone opsin promoter. This is the same flat mount as in Fig. 3A-D. Y-Junctions are denoted by yellow dots, and reverse Y-Junctions are denoted by red dots. **D)** Image of simulation of phase-field crystal model on the surface of a cone. The number of initial rows is comparable to number of initial rows in the retinae. The number of total columns is comparable to the number of total columns in the retinae. The defect density is comparable to the defect density in the retinae. The degree of anisotropy of the triangular lattice is constrained by the anisotropy measured in our live-imaging experiments. Yellow dots denote Y-Junctions. Red dots denote reverse Y-Junctions.

This particle density field is evolved to minimize a free energy, while conserving the total number of particles [60–68] (see Methods). (Although there is no *a priori* reason that the dynamics of a biological system should be governed by a free energy, we expect that any model with symmetric, pairwise, repulsive interactions between cells can be mapped onto an effective free energy.) This free energy incorporates the equilibrium two-point correlation function, which encodes the preferred spacing in the crystal and can be derived from the underlying cone-cone interactions [64–65]. By imaging UV cones near the retinal margin, we know that the two-point correlation function must encode an anisotropic spacing such that the separation between UV cones along the row direction is elongated (Fig. S2).

Given a two-point correlation function, there are only two free parameters within this free energy. The first parameter specifies the degree of undercooling; when the temperature is below the melting temperature (*i.e.*, the liquid is “undercooled”), the density field is unstable to the formation of periodic structures for a range of spatial frequencies. As there is no clear analogue to temperature in our system, we interpret this parameter as quantifying the strength of the interactions relative to the random noise in cone motion. The second parameter is the mean of the cone density field, which is conserved.

The values of these two free parameters, the interaction strength and the mean of the density modulation field, determine which phase (or phases) are stable. Depending on these parameters, this model can generate three phases: a constant (“liquid”) phase, a striped phase, and a triangular phase [60, 66]. We can constrain these two free parameters to the region of the phase diagram in which the only stable phase is the triangular phase. We scan the parameters over a one-dimensional cut of this region of the phase diagram to ensure that our conclusions do not depend on fine-tuning of parameters (Fig. S8; see Methods).

By simulating this model in a geometry of comparable size to the retina and with comparable defect densities, we find that this model generically produces spatial distributions of defects that are quantitatively similar to those observed in the flat-mounted retinae (Figs. 3, 7C-D, S4, S8). The fraction of defects in grain boundaries, as quantified by the measure that we applied to the flat-mounted retinae, is approximately sixty percent (Figs. 3E, S8). In the eight flat-mounted retinae, approximately fifty percent of Y-Junctions are in grain boundaries (Figs. 3E). This model, for which there is supportive evidence in embryonic retinae [43], is consistent with observations of flat-mounted retinae (Fig. 7C-D).

#### Additional insights generated by model of cell-cell repulsive interactions

Armed with this cell-cell repulsion model, we now generate insights into cone mosaic formation. First, we address how the specific orientation of the cone mosaic lattice is selected and how it is maintained as the crystal grows. With isotropic interactions in an isotropic medium, at the onset of ordering, crystallites form with random orientations. Additionally, as the crystal grows outward, the crystallites tend to rotate into an orientation that is misaligned with the orientation observed in the retinae by thirty degrees (*i.e.*, maximally misaligned) (Fig. S9A, Vids. 3-4). For an anisotropic crystal, as in the retinae, the correct crystallographic orientation is selected even without spatial ordering in the original cone positions, and that crystallographic orientation is maintained as the crystal grows (Fig. S9B, Vid. 5).

This model also suggests a mechanism by which grain boundaries form during initial mosaic formation. We consistently observe in our phase-field crystal simulations that the density profiles near a grain boundary remain poorly resolved even as neighboring regions of the crystal grow outward, leading to a characteristic V-shape of the crystal surface (Fig. S9C). Our model predicts that there should be a lag in proper positioning of cones near a grain boundary.

#### Implausibility of lateral inhibition mechanism for cone mosaic formation

Before concluding our modelling of cone mosaic formation, we present a possible alternative model and explain its shortcomings in generating a crystalline mosaic. The alternative model is motivated by other biological lattices like sensory bristles and R8 photoreceptors in *Drosophila* [2–7, 28], where neural cells are selected through a process of lateral inhibition mediated by the Notch signaling system. In these examples, cell motion is largely absent during pattern formation. Mathematical models of motionless tissues in which cells differentiate, inhibiting neighboring cells within some range from committing to the same fate [2-3, 7, 28], reproduce the observed patterns.

The lateral inhibition model that we adapt was originally developed in the context of sensory bristle patterning in *Drosophila* [28] (see Methods). An important difference between sensory bristle patterning in *Drosophila* and cone mosaic formation in zebrafish is that the lattice vectors in the cone mosaic are much shorter, in units of cell diameters, than the lattice vectors of the sensory bristle lattice. There are five to six cells between nearest-neighbor sensory bristle cells [28], but only one to two cells between nearest-neighbor UV cones. Therefore, we expect a cone mosaic lattice generated by lateral inhibition to be more sensitive to disorder in cell packing than is the sensory bristle lattice.

In Figure S10C, we provide an illustrative example of lateral inhibition with a short signaling range on a fixed cell packing (see Methods). As a wave of differentiation moves from the left side of the cell packing to the right, individual cells differentiate into the inhibiting cell fate, and these cells signal to their neighbors, causing them to adopt the alternative fate. The resulting fate pattern has many defects, disrupting the long-ranged crystalline order expected in the cone mosaic [21, 41]. Thus, the disordered cell packing prevents the formation of a precise triangular lattice, in contrast to the previous model in which fate specification occurs first and cells of known spectral fate then move into the correct lattice position.

Since the basic impediment to pattern formation by lateral inhibition in this model is the disorder in the cell packing, one might be tempted to consider a model in which lateral inhibition instead acts to impose a fate pattern on an ordered packing of equipotent cells. In such a model, however, there would have to be defects in the ordered packing to fit onto the retina’s curved surface, and the pattern of defects in the eventual cone mosaic would be expected to follow the pattern of defects in the underlying packing. Thus, the problem would again be reduced to that of arranging cells (necessarily with some short-ranged repulsion, as they cannot overlap) into the surface of a growing hemisphere – that is, essentially to the same problem solved by our earlier phase-field model of cone mosaic formation.

## DISCUSSION

In this study, we characterize the properties of Y-junction defects in the zebrafish cone mosaic and their spatial distribution across the retina. Strikingly, we find that Y-junctions are organized into grain boundaries oriented perpendicular to the retinal margin, as would be expected if they were positioned to minimize the elastic energy of a physical crystal. We show, however, that unlike dislocations in most physical crystals, Y-junction motion is limited, implying that Y-junctions must be positioned within existing grain boundaries when they are created at the cone mosaic’s growing margin. Inspired by these observations, as well as previous findings that cone photoreceptors in embryonic retinae are born from symmetric terminal divisions and then disperse to their final positions in the retina, we propose a model for cone mosaic formation based on interactions between fate-committed cells, generating cell motion. This model reproduces the major features of the Y-Junction distribution in the zebrafish retina.

Our model of cell motion contrasts with most previous pictures of pattern formation in sensory epithelia [2-7, 28, 69-71]. For example, in R8 photoreceptor specification, without invoking cell motion, one reproduces the observed patterns based on cell-cell signaling in a disordered tissue. Cells of an inhibiting fate prevent neighboring cells from adopting the same fate within some signaling range. In both R8 photoreceptor specification and the similar problem of sensory bristle patterning, the signaling range is larger than the typical cell diameter, resulting in approximately six cells between each inhibiting cell [2–7, 28]. These lateral inhibition systems are relatively insensitive to disorder in the cell packing.

We have argued, on the other hand, that if the lateral inhibition signaling range is on the order of the typical cell size in a disordered packing, one must invoke cell rearrangement to produce a crystalline mosaic. This finding is consistent with a very recent study by Cohen *et al.* of the patterning of outer hair cells in the mammalian inner ear [9]. The outer hair cells form a triangular lattice in a quasi-two-dimensional tissue where the lattice spacing is on the order of one cell [8–9]. To produce a pattern consistent with the order observed in the inner ear, Cohen *et al.* first produce a fate pattern via lateral inhibition in a disordered packing. They, then, model rearrangements of the disordered packing: global shear forces and local repulsion between outer hair cells, followed by type-dependent edge tensions. Even with such significant rearrangements, their model produces excess defects in the triangular cell lattice of outer hair cells [9].

In contrast, our model of cone mosaic crystallization is based on interactions between cones of the same subtype (*e.g.*, UV cones) during their differentiation and incorporation into the retina. We hypothesize that these repulsive interactions are mediated by cellular appendages called telodendria [72–73]. Previous investigators have suggested the potential role of telodendria in tangential dispersion after symmetric terminal cell division [43]. Though less ordered than the zebrafish cone mosaic, both the retinal horizontal cell mosaic in mice and class IV neural cell mosaic in *Drosophila* form via similar neurite-mediated interactions, demonstrating the importance of cell motion guided by repulsive interactions for forming neural cell mosaics [44–48].

It is tempting to speculate about the merits of cone mosaic formation via homotypic cell-cell repulsion between fate-specified cones as opposed to a more classic lateral inhibition pathway. For example, if the formation of defect-free crystalline domains (separated by grain boundaries) is functionally relevant, which aspects of fish retinae might allow our model to outperform lateral inhibition on that merit? One possibility is that it is precisely the curvature of the retinal hemisphere and the resulting need for lattice defects that favor the homotypic repulsion mechanism over lateral inhibition. Indeed, we have argued that a cone mosaic formed in such a way will have many of the same features as a physical crystal. These features include, in particular, the presence of an effective long-ranged, elastic interaction between Y-Junctions that is absent in models of fate specification through signaling on a fixed cell packing. Such a long-ranged interaction likely makes it easier to position Y-Junctions correctly across the retina, as exemplified by the spontaneous appearance of grain boundaries in our phase-field crystal model.

Finally, this study highlights the importance of the kinetics of crystal growth in determining the observed spatial distribution of defects. Many studies of physical crystals focus on the agreement between theoretically predicted ground-state defect distribution and experimentally observed defect distributions [24–26, 34] rather than on the kinetics of grain boundary formation. In the zebrafish retinae, we demonstrate the grouping of dislocations into grain boundaries during initial mosaic formation without requiring glide motion. Akin to work by Köhler *et al.* [52], the phase-field crystal simulations of cone mosaic formation suggest the following mechanism by which dislocations are grouped into grain boundaries: a delay of crystal growth near a grain boundary relative to growth of defect-free domains. Biological tests of this prediction await further investigation.

## MATERIALS AND METHODS

### Zebrafish

Fish were maintained at -28C on a 14/10 h light/ dark cycle with standard husbandry procedures [74]. Zebrafish lines, *Tg(-5.5sws1: EGFP)kj9* [75], *Tg(-3.2sws2: mCherry)mi200*7 [21], *Tg(trβ2: tdTomato)* [43], *Tg(-3.2sws2: EGFP)* [76], Tg(*gnat2*:H2A-CFP), and pigment mutant *ruby* carrying *albino (slc45a)^b4/b4^* and *roy^a9/a9^* were used. All animal procedures were approved by the Institutional Animal Care and Use Committee at the University of Michigan.

### Histology

Retinal dissection, fixation and immunocytochemistry were performed as previously described [41, 77–78]. Briefly, the isolated retina was fixed in 4% paraformaldehyde with 5% sucrose in 0.1M phosphate buffer, pH 7.4, at 4C overnight. After antigen retrieval with 10 mM sodium citrate in 0.05% tween 20 (pH 6.0), retinas were incubated in blocking buffer for 2 hours followed by primary antibody incubation, mouse anti-Zonula Occludens (ZO1-1A1, 1:200, ThermoFisher Scientific, Waltham, MA) and rabbit anti-GFP (1:200, ThermoFisher Scientific) at room temperature overnight. Incubation with secondary antibodies (Alexa Fluor 555 and 649, ThermoFisher) were performed at room temperature overnight, and the retina was flat-mounted on a glass slide. For retinal cross sections, affinity-purified rabbit polyclonal opsin antibodies, a gift from Dr. David R. Hyde [79], were used. Images were acquired with a Zeiss AxioImage ZI Epifluorescent Microscope (Carl Zeiss Microimaging, Thornwood, NY) equipped with an ApoTome attachment for optical sectioning structured illumination, Leica DM6000 Upright Microscope System (Leica Microsystems, Werzlarm Germany) and a Leica TCS SP5 confocal microscope equipped with Leica 40X HCX PL APO CS Oil Immersion lens.

### Generation of transgenic zebrafish with nuclear-localized, photoconvertible (green-to-red) EOS protein expressed specifically in UV cones

Multi-Gateway-based *tol2* kit system was used to generate expression vectors [80]. In brief, the 5’ entry clone, *p5E*-*5.5sws1* [75], middle entry clone, *pME-nEOS* (gift from Dr. David Raible), and 3’ Entry clone, *p3E-polyA*, were assembled into a destination vector, *pDestTol2pA* [80] using LR Clonase II Plus enzyme (Thermo Fisher Scientific). Embryos of the transparent *ruby* genetic background [41] at the 1-cell stage were injected with 1 nL of solution containing 25 pg plasmid DNA and 25 pg tol2 transposase mRNA [81]. Founders (F0) with germline transmission of the transgene were identified by outcrossing with wildtype animals, and their F1 progenies were screened for nEOS expression at 4 days post fertilization.

### CRISPR-Cas9 mediated mutation in the *thrb* gene

A genetic mutation targeting the type 2 isoform of the *thrb* gene (synonym, *trß2*) was generated by CRISPR-Cas9 gene editing methods [82]. Briefly, pT7 gRNA vector (Addgene #46759) was used as a template to construct the *thrb2* gRNA [82]. PCR based method was performed using specific primers, 5’-GGGGTAATACGACTCACTATA GGCAACACAGCCAACCCTATGTTTTAGAGCTAGAAATAGCAAG-3’; 5’-AAAAAGCACCGACTCGGTGCC-3’. The MEGAscript T7 Kit (Ambion Inc., Austin, TX) was used to transcribe the gRNA. For the *nlsCas9nl* mRNA synthesis and purification, mMESSAGE mMACHINE T7 Transcription Kit (Ambion) and Qiagen RNeasy mini kit (Qiagen, Hilden, Germany) were used. For genotyping of the *thrb* mutation, PCR fragments of the *thrb* gene, amplified using specific primer set, 5’-CATGGTGTAAGTGGCGGATATG -3’; 5’-TCCACTGCATCTGAGAGAAATCC-3’, were subjected to restriction with BstXI (New England Biolabs, Ipswich, MA).

### nEOS photoconversion and imaging

Photoconversion of nEos protein was performed on *ruby; Tg(sws1:nEos)* fish [83]. Juvenile zebrafish (0.7 to 0.88 cm standard body length) were anesthetized with 0.672 mg/ml Tricaine S/ MS-222 (Western Chemical Inc., Ferndale, WA) and placed dorsal side down on a 50 mm glass bottom petri dish with a No. 1.5 coverslip (MarTek Corporation, Ashlan MA, see [41]) and held in place with damped Kimwipes. Imaging and photoconversion were performed with a Leica TCS SP8 LSCM (Leica Microsystems, Werzlar, Germany) equipped with Leica 40X PL APO CS2 Water Immersion lens, 1.1 NA with 650 μm working distance. Green to red photoconversion of nEOS protein was performed by a 405 Diode laser at 400 Hz scan speed with a resolution of 512 x 512 pixels in the *xy* dimension at a single optical plane. Pre and post photoconversion images were captured with the White Light Laser tuned to 506 nm for nEOS (green) and 573 nm for nEOS (red). Leica HyD hybrid detectors were tuned to 516-525 nm for nEOS (green) and 620-761 nm nEOS (red).

### Large tile scans of flat-mounted retinae

Large tile scans of entire flat-mounted retinae from adult *Tg(sws1:EGFP)* zebrafish immunostained for ZO1 were acquired with a Leica TCS SP8 LSCM (Leica Microsystems) equipped with Leica 20X PL APO Dry lens. The GFP signal was recovered by immunostaining with anti-GFP antibody. The White Light Laser was tuned to 555 nm for Alexa Fluor 555 and 649 nm for Alexa Fluor 649. The Leica HyD hybrid detectors were tuned to 600-641 nm for Alexa Fluor 555 and 701-751 nm for Alexa Fluor 649. Images were acquired at 700 Hz scan speed with a resolution of 2048 x 2048 pixels in the *xy* dimension with a μm interval between optical sections in the z-dimension.

### Row tracing of flat-mounted retinae

We manually traced rows of UV cones, starting near the region coinciding with the disorder-to-order transition (Figs. 3A-E, S4; Table S1). The row tracing extends over approximately one hundred columns of UV cones from the larval remnant to the periphery, avoiding regions of the retinae which were damaged during flat-mounting. Based on the row-tracing, we identified where rows are inserted (*i.e.*, Y-Junctions) and where rows are removed (*i.e.*, reverse Y-Junctions, see Table S1).

### Detection of Grain Boundaries

In the flat-mounted retina, we have the positions of all Y-Junctions, which generate row insertions, in the traced regions. To define grain boundaries, we search for approximately linear chains of five Y-Junctions, which are nearest neighbors. To the image of all Y-Junctions in the traced regions (Fig. S4), we apply the following algorithm:

1. Loop through all Y-Junctions one-by-one. We will build a chain of nearest neighbors, of five Y-Junctions, for each Y-Junction.

a. Look for the Y-Junction’s nearest neighboring Y-Junction, using a k-nearest neighbors search (knnsearch; MATLAB 2016B, MathWorks, Natick, MA). Add the nearest neighbor to the chain.
b. For that nearest neighbor, add its nearest neighbor, excluding any Y-Junctions which already belong to the chain.
c. Repeat b until you have a chain of five Y-Junctions, including the Y-Junction which initialized the chain.
2. Now, based on the calculation in step 1, every Y-Junction, indexed by *i* below, has a chain of five nearest neighbors, including the Y-Junction itself. We index the five defects in the chain by *j*. The position of the *j*^*th*^ defect in the *i*^*th*^ chain is 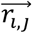. For each Y-Junction, we compute the following sum:

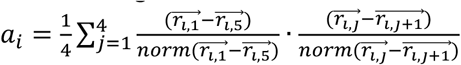
3. Now, based on the calculations in steps 1 and 2, every Y-Junction, indexed by *i*, has a chain of five nearest neighbors, which is assigned a score *a*_*i*_. If *a*_*i*_ = 1, the chain of five Y-Junctions is perfectly linear. If *a*_*i*_ > *a*_*gb*_, we call the *i*^*th*^ chain a grain boundary. We set the cutoff *a*_*gb*_ equal to seven-eighths.
4. We initialize an empty array, in which we will store Y-Junctions which belong to grain boundaries. Loop through all Y-Junctions, indexed by *i*, one-by-one. If *a*_*i*_ > *a*_*gb*_, store (i.e., in the array of all Y-Junctions in grain boundaries) the five Y-Junctions which belong to this chain.

We recognize a couple of limitations of this approach as applied to the flat-mounted retina. First, for flattening the retina, the retina is sliced in four places. The cuts could physically separate one part of a grain boundary from another, leading us not to detect the grain boundary. Second, the flattening deforms the cone mosaic, which means that spatial relationship between Y-Junctions is somewhat different from the intact hemispheric retinae. Nevertheless, the algorithm performs well in identifying grain boundaries in the retina, as illustrated on three of the flat-mounted retinae in Fig. S4.

We also perform this computation (steps 1-4) on the seven-coordinated particles in simulations of the phase-field crystal model. In that case, we map the cone frustum to the flat plane. This does not generate distortions because cones are isometric to the plane. We, then, calculate which dislocations form grain boundaries, respecting the periodic boundary conditions of the flattened cone frustum.

### Tracking Nuclear Positions in Photoconverted Regions

In the photoconversion experiments, we observe the same region of the same retina at two different times in live fish. Given a nucleus at one time point, we want to find the same nucleus in the image at the other time point. One image of the region is taken immediately after photoconversion, which we call day 0. Across fish, we vary the time between photoconversion and the time of the second observation (i.e., two days after photoconversion at the earliest and four days after photoconversion at the latest). We call the second time point day 2-4.

At both times of observations for each fish, we have an image with two channels. One channel corresponds to the color of the photoconverted fluorescent protein. The other channel corresponds to the color of the non-photoconverted fluorescent protein. For the image analysis below, we use the photoconverted channel at both times. The image is three-dimensional, and the plane which contains the UV cone nuclei (i.e., where the fluorescent protein is localized) is mostly parallel to the x-y plane. This fact allows us to perform most of the computations, for tracking each nucleus from one image to the other, based on two-dimensional projections.

For each z-stack, we compute a two-dimensional wiener filter (wiener2; MATLAB 2016B Image Processing Toolbox, MathWorks) with a filter size of eight pixels, which is approximately a micron. This filter removes noisy specks (i.e., spikes in intensity at small length scales). We, then, compute a two-dimensional projection by summing over z-stacks. The photoconverted UV cones are in the middle of the image. The intensity in the photoconverted channel is significantly weaker for UV cones near the edge of the image. This provides us the reference boundary by which we can identify common nuclei (i.e., which nucleus in the day 2-4 image corresponds to a specific nucleus in the day 0 image).

We perform an image registration, computing the combination of rotation and translation which optimizes the normalized cross-correlation between the two images (normxcorr2; MATLAB 2016B Image Processing Toolbox, MathWorks). Then, we segment nuclei in the two images. Because the intensity of UV cone nuclei varies significantly across the image, we use both adaptive thresholding (adaptthresh; MATLAB 2016B Image Processing Toolbox, MathWorks) and a low absolute threshold. We morphologically open the thresholded image, followed by morphological closing. We fill holes in the image (imfill; MATLAB 2016B Image Processing Toolbox, MathWorks) and clear the border of the image (imclearborder; MATLAB 2016B Image Processing Toolbox, MathWorks). We perform minimal manual correction of these segmentations. Given that we have aligned the two images and segmented the nuclei, we track each nucleus from one image to the other by computing for each nucleus in the day 0 image its nearest neighbor in the day 2-4 image (knnsearch; MATLAB 2016B, MathWorks). As a sanity check, for each nucleus in the day 2-4 we compute its nearest neighbor in the day 0 image to make sure that calculation returns the same answer for each nucleus. We manually correct any errors.

Following this segmentation and identification of common nuclei between the two images, we want to estimate the three-dimensional position of each nucleus based on the raw z-stacks rather than on a post-processed version. We identify a circular region, of radius two and a half microns, in the xy-plane centered on each of the segmented nuclei. This radius is larger in the xy-plane than the nuclear radius but small enough not to encompass other nuclei. This circular region corresponds to a pillar in the z-direction. To estimate the three-dimensional position of each nucleus in both images, we use the raw z-stacks, computing the center of intensity of each pillar (i.e., weighted average of voxel positions in each pillar where the weights are the voxel intensities). At the end of this entire procedure, for each nucleus common to both images, we know its position at both time points.

### Triangulating UV Cone Positions in Photoconverted Regions and Measuring Glide Motion

To identify the location of the Y-Junction, we need to calculate a triangulation over the nuclear positions. At both day 0 and at day 2-4, the UV cone nuclei positions in each experiment are well fit by a plane, which we fit by simple least-squares minimization (see RMSE information in Table S2). For calculating the triangulation, we project the UV cone positions onto the plane of best fit. We, then, calculate the triangulation in that plane (delaunayTriangulation; MATLAB 2016B, MathWorks).

We want to track movement of UV cones near the Y-Junction core along the direction of glide motion. We systematically search for bond flips (*i.e.,* any change in nearest neighbor assignments as in Fig. 5C) between day 0 and day 2-4 for any bonds that could be flipped in glide motion (see Fig. 4C). Which UV cone bonds lie along the glide line is **always** unambiguous based on the triangulation. We never observe glide motion by more than one unit, as illustrated in Figs. 4C, 5A. We show an experimental example of glide motion by one unit in Fig. 5C (see Table S2).

### Statistical Significance of Growing Grain Boundaries in Live Fish

In Figure S6, we show examples of images in which we can identify newly incorporated Y-Junctions lining up into grain boundaries. These are images of UV cone nuclei near the retinal margin (*i.e.*, where the layer grows by addition of post-mitotic cells). These images are oriented such that the margin is parallel to the y-axis. Because our field of view in these images is limited (*i.e.*, approximately forty rows of UV cones and forty columns of UV cones), it is difficult to apply our grain boundary measure that we use for flat-mounted retinae and simulations (see Detection of Grain Boundaries in Methods).

To identify grain boundaries which are already visible immediately after photoconversion, we trace rows of UV cones at the retinal margin. If immediately after photoconversion the row direction rotates about a group of defects by ten degrees or more at the margin, we call the group of Y-Junctions in-between the rotated rows a grain boundary. As a justification for this use of row rotation to identify grain boundaries, see Figs. 3C-D, 5D, S6. Based on this criterion, out of the eighteen samples, twelve samples have grain boundaries near the retinal margin at the time of photoconversion. Since some samples have two grain boundaries, in total we observe fifteen grain boundaries (Fig. 5E).

All subsequent analysis is based on the later image (*i.e.,* two, three, or four days later). We trace rows in the later image. We identify newly inserted rows (*i.e.,* newly incorporated Y-Junctions) in the later image, and we again identify the old defects within each grain boundary (*i.e.,* those not newly incorporated). We calculate a one-dimensional coordinate for the location of each grain boundary in the later image. This one-dimensional coordinate is the average of y-coordinates (*i.e.*, axis approximately parallel to the margin in the image) of all defects (*i.e.,* not newly incorporated) within each grain boundary. We are interested in how close, along the y-direction, newly incorporated Y-Junctions are to the nearest grain boundary in the image.

Suppose there is only one grain boundary in the image which is identifiable at the time of photoconversion and later imaging. Suppose this grain boundary is located at coordinate *y*_*gb*_. The image spans from *y* = 0 to *y* = *y*_*max*_. For each new Y-Junction, we generate one hundred thousand random Y-Junction positions, uniformly distributed from *y* = 0 to *y* = *y*_*max*_ (rand; MATLAB 2016B, MathWorks). We calculate the distance between each of these one hundred thousand random Y-Junction positions and the grain boundary (at *y*_*gb*_). We call the vector of distances between each random Y-Junction position and the grain boundary 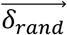 We also store the actual distance, which we call *δ*_*actual*_, between each observed newly incorporated Y-Junction position in the image and the nearest grain boundary in the image.

Suppose there are two grain boundaries in the image which are identifiable at the time of photoconversion and later imaging. Suppose their coordinates are *y*_*gb*,1_ and *y*_*gb*,2_. The image spans from *y* = 0 to *y* = *y*_*max*_. For each new Y-Junction, we generate one hundred thousand random Y-Junction positions, uniformly distributed from *y* = 0 to *y* = *y*_*max*_ (rand; MATLAB 2016B, MathWorks).

We calculate the distance between each of these one hundred thousand random Y-Junction positions and the nearest grain boundary (at either *y*_*gb*,1_ or *y*_*gb*,2_). We call the vector of distances between each random Y-Junction position and the nearest grain boundary 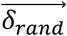 We also store the actual distance, *δ*_*actual*_, between each newly incorporated Y-Junction position and its nearest grain boundary.

Based on the procedure outlined above, for each new Y-Junction, we have a vector of length one hundred thousand and a scalar, 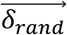 and *δ*_*actual*_(see Table S3). If after random incorporation with respect to the grain boundaries in the image, a newly incorporated Y-Junction moves at a speed of one row per two days closer to the nearest grain boundary (with spacing between rows approximately equal to six microns as shown in Fig. S2), the distribution of distances with respect to the nearest grain boundary becomes max 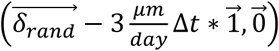, where Δ*t* is the time between photoconversion and later imaging (*i.e.,* two, three, or four days).

We have a total of thirty-seven new Y-Junctions across the twelve samples with grain boundary at the retinal margin. We would like to compare the thirty-seven scalar values of *δ*_*actual*_ to a concatenated vector of max 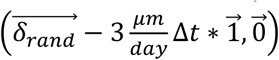 across all thirty-seven defects. This concatenated vector is of length three million seven hundred thousand. We test whether the distribution of *δ*_*actual*_ has the same median as the concatenated vector of max 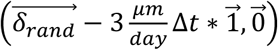 across all thirty-seven defects. We assign a p-value to that comparison via Mann-Whitney U-test (ranksum; MATLAB 2016B Statistics and Machine Learning Toolbox, MathWorks).

### Numerical Solutions of Lateral Inhibition on Disordered Cell Packing

Starting with a Voronoi tessellation of uniformly (randomly) distributed points, we generated large, disordered, periodic cell packings (*e.g.*, 20,000 total cells in Fig. S10) via vertex model simulations with equal tensions on all edges as described in [84–85]. We model dynamics of individual cell fates on the static cell packing according to the model described in [28], but do not include noise in the dynamics (*D* = 0). Since we changed some aspects of the model, including the external signaling gradient and the noise in fate, we describe the model in [28] below for the sake of clarity. The fate of cell *i*, *u*_*i*_, evolves as:

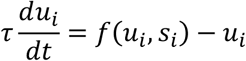

where *s*_*i*_ is the signal each cell receives from other cells as well as from any external gradients. We interpret the *u* = 1 fate as the UV cone spectral subtype and the *u* = 0 fate to be other spectral subtypes. Also, *f*(*u,s*) is sigmoidal: 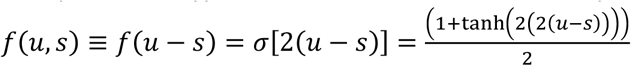.

The signal that cell *i* receives, *s*_*i*_, includes an external time-dependent signal *s*_0_(*x*, *t*) as well as signals from neighboring cells in a distance-dependent manner.

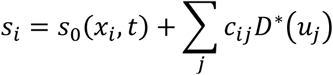

The external signal provided to the cells has the following form: 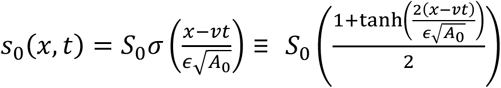 where *S*_0_ = 1 and 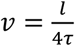 and 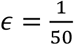. *τ* is the timescale for cell fate dynamics, and *l* is the characteristic cell-cell signaling range. The distance-dependent coupling constant *c*_*ij*_ between cell *i* and cell *j* is of the form: *c_ij_* = e^-*dij*2/(2*l*2)^, where *d_ij_* is the distance between the centroids of cell *i* and cell *j*. No cell signals to itself directly: *c*_*ii*_ = 0.

A cell of fate *u*_*j*_ produces signal *D*^∗^(*u*_*j*_) = *a*(*u*_*j*_)*D*(*u*_*j*_). The ligand level of cell *j*, *D*(*u*_*j*_), is directly proportional to the fate *u*_*j*_. The ligand activity of cell *j*, called *a*(*u*_*j*_), is of the form: 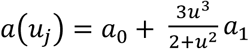 *a*_1_. We use the same ligand activity levels for both the *u* = 0 fate and the *u* = 1 fate as in [28] (*a*_0_ = 0.05; *a*_1_ = 1 − *a*_0_).

To explore the effects of cell-cell signaling range on the final fate pattern, we systemically change the signaling range *l* from 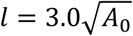 to 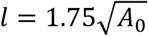 to 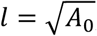, where *A*_0_is the mean cell area. All cells are initially in the *u* = 0 state. The sigmoidal signaling front, sharper than the characteristic cell size, starts at left side of the packing (*x* = 0 at *t* = 0) and moves to the right. In the wake of the front, individual cells differentiate into the *u* = 1 fate, inhibiting their neighbors from adopting the *u* = 1 fate within the specified cell-cell signaling range. We solve the differential equations for cell fates using ode45 (MATLAB 2016B, MathWorks).

### Numerical Solutions of Anisotropic Phase Field Crystal Model on Cone

The free energy *F* for an anisotropic phase field *ψ* [60, 64–65]:

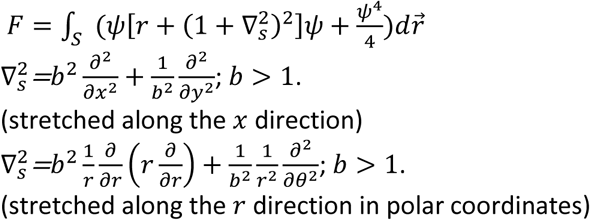

The particle density modulation field *ψ* is evolved to minimize the free energy *F* while conserving the 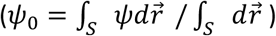

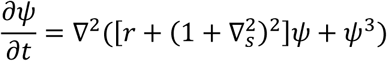

For solving this equation on the cone, we first map the cone to a flat plane, which does not generate distortions because the cone is isometric to the plane. We, then, use the Laplacian for polar coordinates, respecting the periodic boundary condition of the cone. We set up in the problem in terms of the variables *u*, *v*, *ψ* as defined in [86]. For computational efficiency, we take the Fourier transform along any direction which is periodic (e.g., along the *θ* direction on the cone). We use first-order implicit-explicit methods, as in [87], treating the non-linear term in *ψ* explicitly. We implement all derivatives by finite differences [88]. We use no-flux boundary conditions at each non-periodic boundary.

We evolve the system with a fixed step size in time (Δ*t* = 0.075). The computational grid is such that there are approximately 25 grid points per lattice spacing along the circumferential direction in the initial row, and approximately 10 grid points per lattice spacing along the radial direction.

We systematically vary the parameters of the phase field crystal model as done in [60]; please note that stretching the crystal does not change the phase diagram as discussed in [66]. In short, we take a 1D cut of the phase diagram, setting 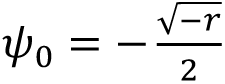 as we vary the undercooling parameter *r*. For this cut, we also vary the strength of the noise in the initial conditions These three parameters are the parameters on which we do not have any quantitative handle (relative to experiments); therefore, we perform this robustness analysis on these three parameters. The results of this analysis are shown in Fig. S8.

#### Geometry for Cone Mosaic Growth

The retina is approximately hemispheric. A hemisphere might, thus, seem like the most obvious choice of geometry in which to test cone mosaic growth. It is important to note, however, that the retina is not a hemisphere of a fixed radius during development. Its radius increases as new retinal cells are incorporated. As the hemisphere dilates, the existing cone photoreceptor layer must be deformed. The exact way in which the existing pattern deforms, and how that affects subsequent cone mosaic formation, is beyond the scope of this paper. Our aim is to choose a minimal geometry which allows us to test the phase-field crystal model’s ability to form grain boundaries.

We choose a geometry in which we can easily tune the defect density. We choose a cone frustum, which is constructed by slicing off the top of a cone with a plane that is parallel to its base. By changing the level at which we slice the cone, we tune the number of UV cones in the initial column. We choose the top level such that there are approximately two hundred initial rows, which is consistent with the number of initial rows identified in flat-mounted retinae. By changing the opening angle of the cone, we can tune the number of Y-Junctions required to maintain constant cell-cell spacing per added column. We choose an opening angle such that two row insertions are required per added column, to maintain approximately constant cell spacing. The number of Y-Junctions necessary to maintain constant cell-cell spacing is comparable to the number of Y-Junctions observed in the retinae (Figs. 3E, S4).

#### Initial Conditions for Cone Mosaic Growth

At the very top level of the cone frustum, we lay down one column of cones (see one-mode approximation in [60]). We add a white-noise mask to this initial column of cones. Because we do not have any quantitative handle on the noise in cell positions at the onset of patterning in the zebrafish retina, we vary the noise strength, exploring its effect on the subsequent pattern of defects (Fig. S8).

## ACKNOWLEDGEMENTS

Linda K. Barthel provided expert technical assistance with confocal microscopy for large tile scanning and photoconversion. We thank David R. Hyde for the gift of opsin antibody and David Raible for the gift of the nEOS Gateway clone. Undergraduate students Komal Govril and Gene R. Bell assisted with tracing UV cones in retinal flat mounts.

## FUNDING SOURCES

We acknowledge funding from the following sources: National Science Foundation IOS 1353914, National Institutes of Health (NEI) P30 EY07003, and National Science Foundation DGE 1256260.

## VIDEO CAPTIONS

**Video 1. Formation of anisotropic triangular lattice in a rectangular domain.** The top and bottom (y) boundaries are periodic; the left and right (x) boundaries are no-flux. There are one hundred rows and one hundred columns.

**Video 2. Formation of anisotropic triangular lattice on a cone frustum with one initial column (masked with white noise).** The cone is mapped to a flat plane here. The inner boundary (closest to origin) and the outer boundary (farthest from origin) are no-flux boundaries. The remaining two boundaries are periodic. There are two hundred forty initial rows and approximately one hundred sixty columns.

**Video 3. Formation of isotropic triangular lattice on a cone frustum with one initial column (masked with white noise).** The cone is mapped to a flat plane here. The inner boundary (closest to origin) and the outer boundary (farthest from origin) are no-flux boundaries. The remaining two boundaries are periodic. There are two hundred forty initial rows and approximately one hundred sixty columns. Note that the dimensions of this simulation domain are different than that of video 2 (because this lattice has isotropic UV spacing, not anisotropic UV spacing).

**Video 4. Formation of isotropic triangular lattice on a cone frustum with only white noise mask (no initial column prepattern).** The cone is mapped to a flat plane here. The inner boundary (closest to origin) and the outer boundary (farthest from origin) are no-flux boundaries. The remaining two boundaries are periodic. The dimensions of the simulation domain are the same as in video 3.

**Video 5. Formation of anisotropic triangular lattice on a cone frustum with only white noise mask (no initial column prepattern).** The cone is mapped to a flat plane here. The inner boundary (closest to origin) and the outer boundary (farthest from origin) are no-flux boundaries. The remaining two boundaries are periodic. The dimensions of the simulation domain are the same as in video 2.

## SUPPLEMENTARY FIGURES

**Supplementary Figure 1.**
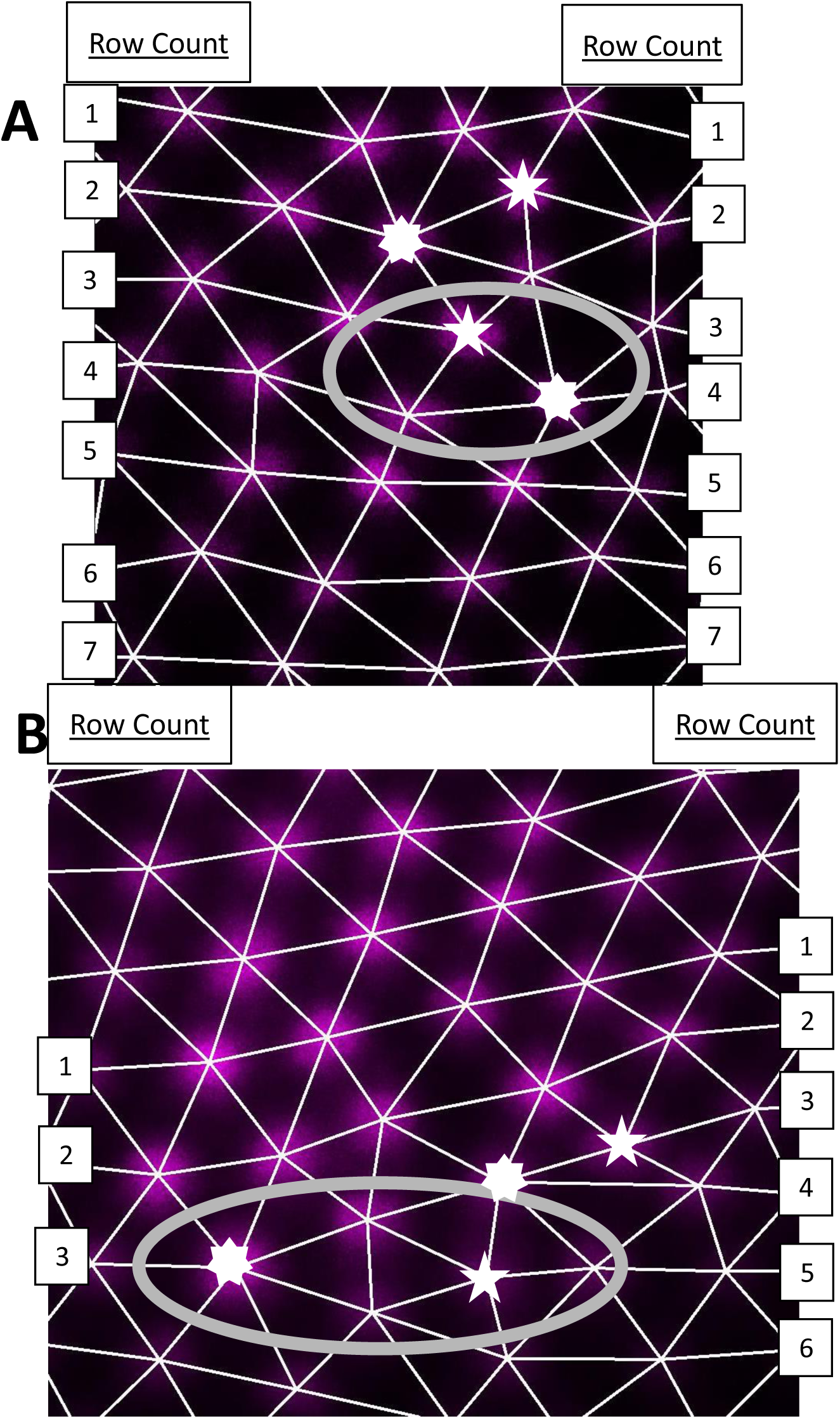
Defects other than Y-Junctions observed in live fish. **A)** In this image, the nuclear-localized, photoconverted protein in UV cones is pseudo-colored magenta. The triangulation that connects nearest neighbors in the lattice is denoted by the white-bonds. The seven- and five-fold coordinated UV cones are denoted by seven- and five-sided stars, respectively. Note that there is a standard Y-Junction near the reverse Y-Junction. The reverse Y-Junction is enclosed by the gray oval. Row counts are annotated on each side of the image. **B)** In this image, there exists both a double-row insertion and a standard Y-Junction. The double-row insertion, enclosed by the gray oval, corresponds to a five- and seven-coordinated particle which are not directly connected by a bond in the lattice. Note that this double-row insertion does not disrupt the patterning of the cone mosaic. Row counts are annotated on each side of the image.

**Supplementary Figure 2.**
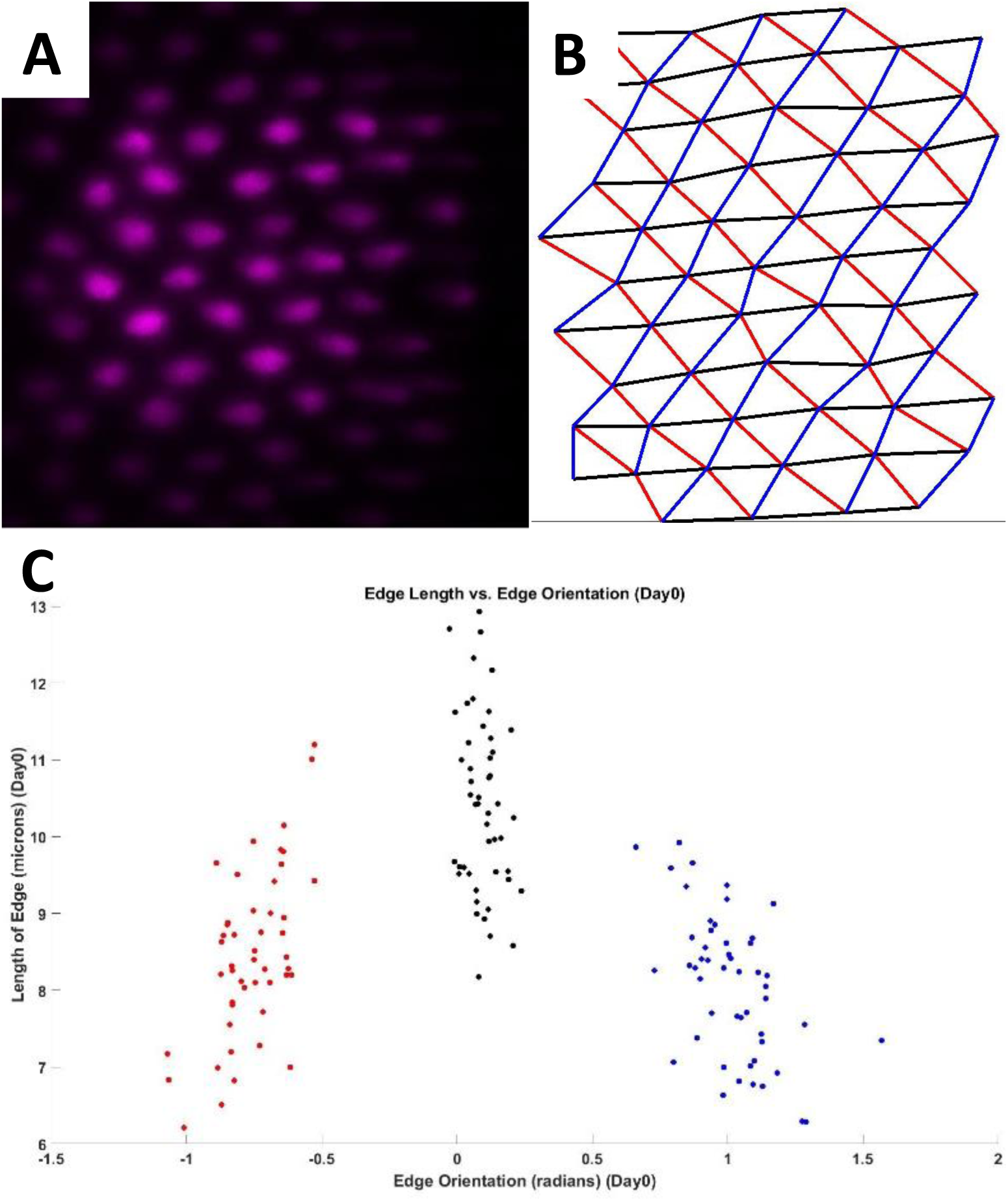
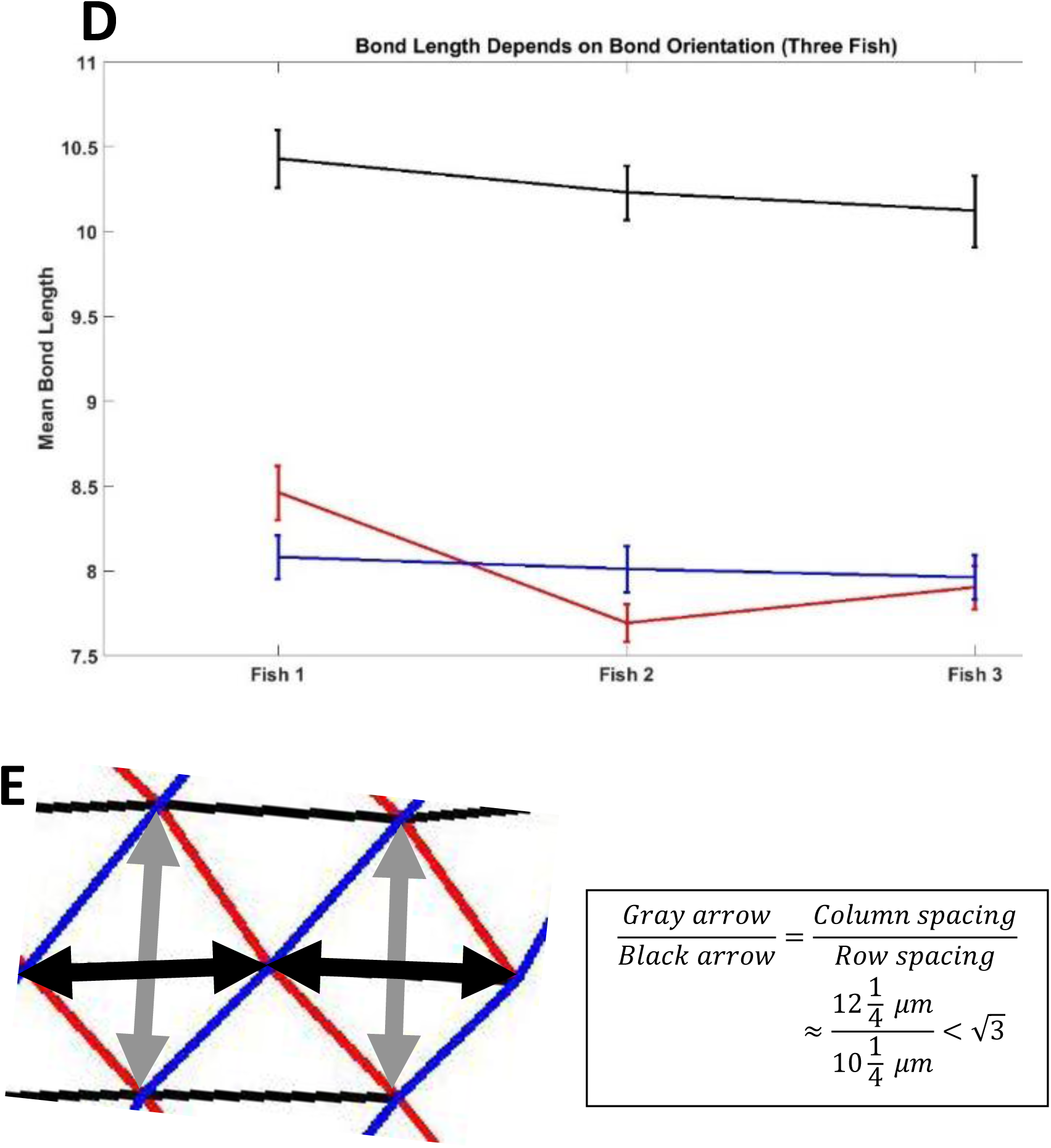
In the live-imaging experiments of UV cones, we can quantify the anisotropy of the UV cone triangular lattice. **A)** A patch of photoconverted UV cones near the retinal margin. This patch of UV cones, which express a nuclear-localized fluorescent protein, does not contain a Y-Junction. We use this patch, and two others like it, to quantify the spacing between UV cones in the mosaic. B) The triangulation that corresponds to the patch of UV cones in panel A. In this triangulation, bonds that connect UV cones in the same row are denoted by black lines. Bonds along the other two principal directions in the triangular lattice are denoted by blue and red bonds. C) Scatter plot of the bond length vs. bond orientation in the triangulation from panel B. The same color scheme is used to denote bonds which are along the row direction and along the two other principal directions. D) For the three photoconverted patches in which there were no Y-junctions, we calculated the mean bond length (and standard deviation of the mean bond length) along the three principal directions. The same color-scheme is employed such that the black points correspond to the mean bond length along the row direction, and the red and blue points correspond to the mean bond length along the other two principal directions in the triangular lattice. E) The column direction is NOT a principal direction in the triangular lattice, meaning that UV cones in the same column are not each other’s nearest neighbors in the lattice. Using a section of the triangulation from panel B, we illustrate the spacing along the row direction by the black arrows, and the spacing along the column direction by the gray arrows. For an isotropic lattice, the column spacing is a square root of three times the row spacing. For this lattice, we can calculate the column spacing, given the mean bond lengths in the three principal directions. Using an identity for the height of an isosceles triangle, we find that the column spacing is approximately twelve and a quarter microns, as compared to a row spacing of approximately ten and a quarter microns. This column spacing to row spacing ratio is less than a square root of three, which means that the row bonds are elongated in comparison to an isotropic triangular lattice.

**Supplementary Figure 3.**
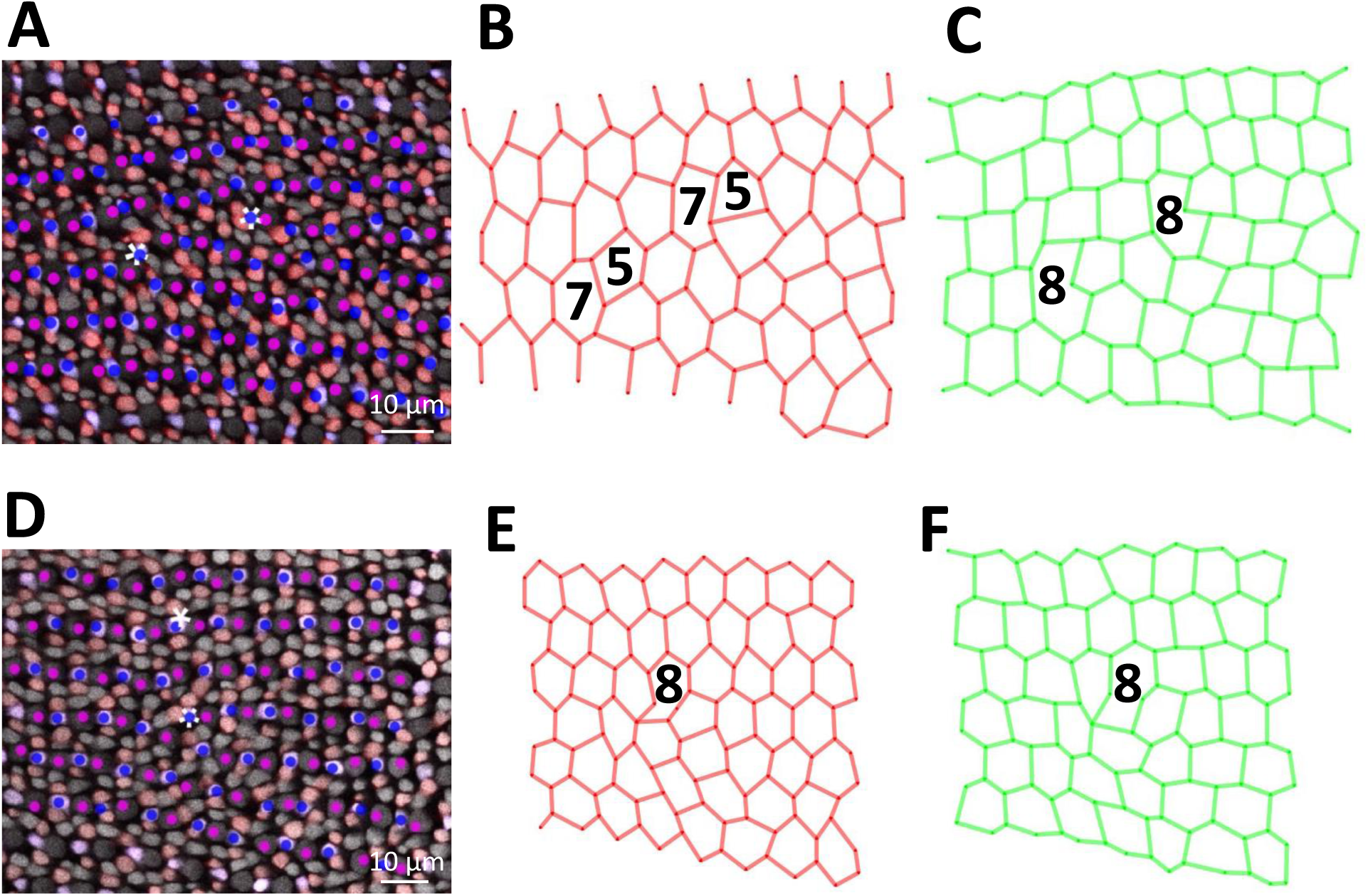
Characterization of the distribution of Red and Green cones near the core of a Y-Junction. **A)** Two Y-Junctions (asterisks) in a flat-mount retinal preparation from an adult, triple transgenic (Tg[*sws2:GFP*; *trβ2:tdTomato*; *gnat2:CFP*] line) in which Blue cones express a fluorescent reporter (pseudo-colored blue) under the control of the Blue opsin promoter *sws2* and in which Red cones express a fluorescent reporter (pseudo-colored red) under the control of the *trβ2* promoter. All cones express an additional fluorescent reporter under the control of the *gnat2* promoter. Please note that although UV and Green cones are not expressing different fluorescent reporters, one is able to distinguish between these two cone subtypes based on morphological differences. B) The nodes of the graph correspond to Red cones from panel A, and the edges connect nearest neighbors in this honeycomb lattice. Note the existence of a heptagon-pentagon pair (*i.e.*, a ‘glide’ dislocation) in the core of both defects. C) The nodes of this graph correspond to Green cones from panel A, and the edges connect nearest neighbors in the honeycomb lattice. Note the existence of an octagon (*i.e.*, a ‘shuffle’ dislocation) in the core of both defects. D-F) Another example of a Y-Junction from a flat-mount retinal preparation from the same double transgenic line (akin to panels A-C).

**Supplementary Figure 4.**
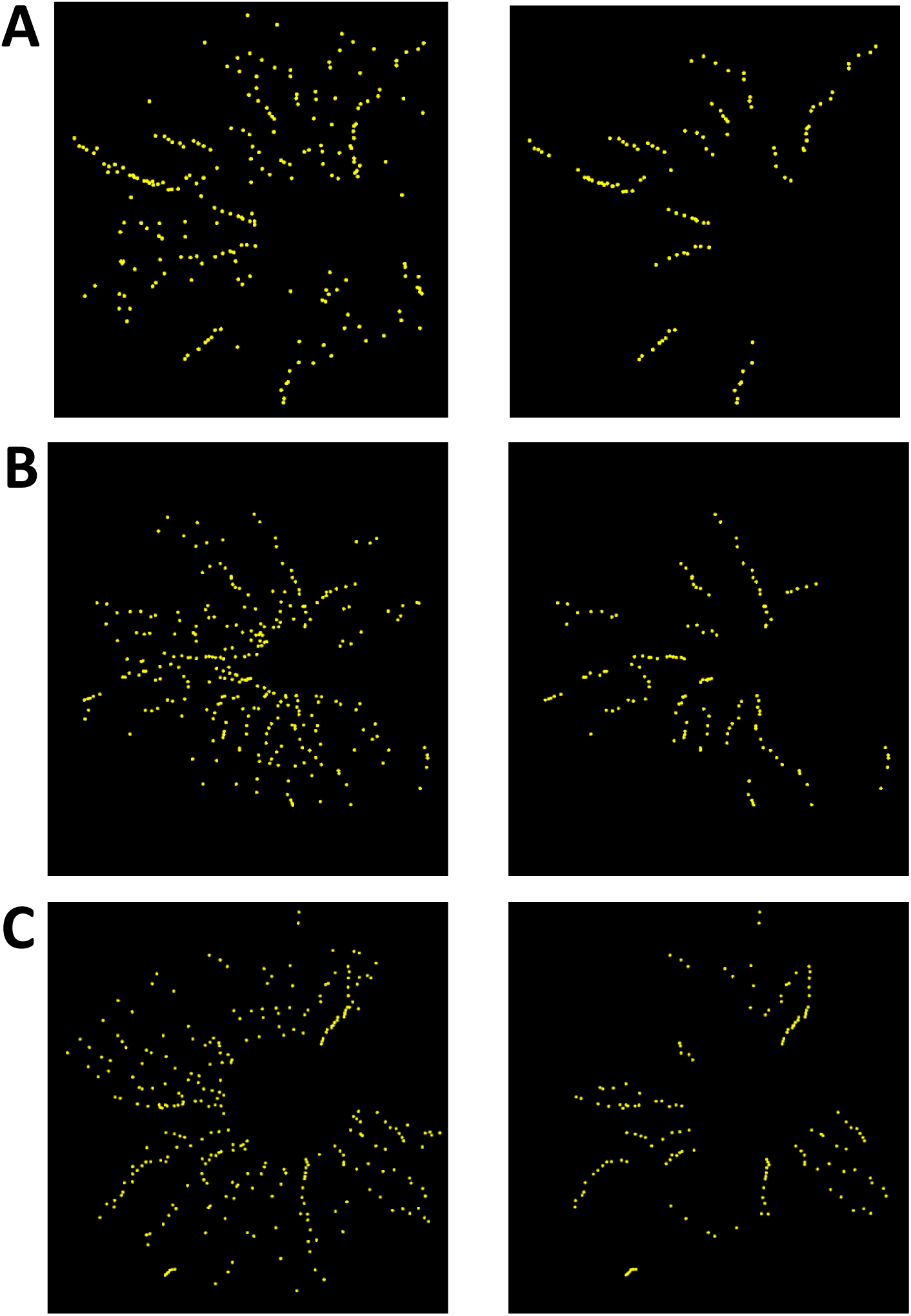
Application of our algorithm for identification of grain boundaries to three flat-mounted retinae. **A-C)** For each of three flat-mounted retinae, the image on the left-hand side is the image of all identified Y-junctions, denoted by yellow dots. The image on the right-hand side is the image of all Y-junctions which our measure determined to be inside of a grain boundary. For each of the three fish, the Y-Junctions in the right-hand image are a subset of the Y-Junctions in the left-hand image. Panel A is fish 3 (total number of Y-Junctions = 221; number of Y-Junctions in grain boundaries = 105). Panel B is fish 4 (total number of Y-Junctions = 275; number of Y-Junctions in grain boundaries = 132). Panel C is fish 8 (total number of Y-Junctions = 285; number of Y-Junctions in grain boundaries = 144), the retina in Figs. 3, 6.

**Supplementary Figure 5.**
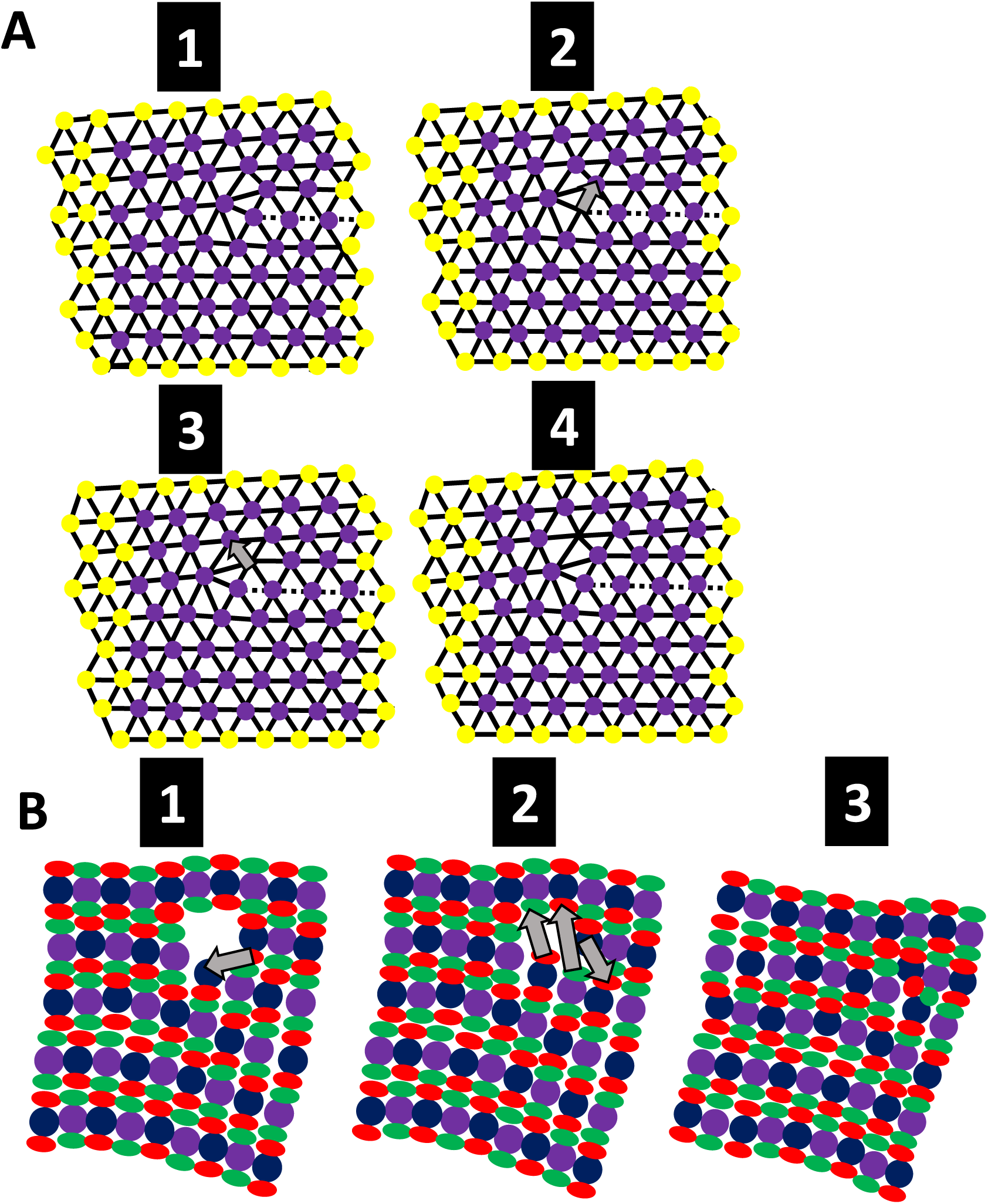
Climb motion requires the creation or annihilation of vacancies or interstitials, which have no analog in the cone mosaic. **A)** The creation of a vacancy allows dislocation to climb (i.e., move perpendicular to Burgers vector). The lattice in panel 1 corresponds to a lattice with a dislocation with photoconverted UV cones in magenta, and non-photoconverted UV cones in yellow. Panel 2 is the triangulation in panel 1 with a new vacancy. The gray arrow denotes where the vacancy will hop, to create the distribution in Panel 3. The gray arrow in Panel 3 denotes where the vacancy will hop, to create the distribution in Panel 4. Note that as the vacancy hops away from the dislocation core, the defect core moves (i.e., perpendicular to the Burgers vector). B) A vacancy in the cone mosaic (which involves two missing Red cones, two missing Green cones, one missing Blue cone, and one missing UV cone) can be destroyed, which allows for movement of the defect core. The Red cone in Panel 1 must move as indicated by the gray arrow to create the distribution in Panel 2. The movements denoted by the gray arrows in Panel 2 allow for the vacancy to close, and for the defect to move. Panel 3 corresponds to the distribution of cones after the vacancy has been destroyed. We never observe such a vacancy (involving two missing Red cones, two missing Green cones, one missing Blue cone, and one missing UV cone) in the cone mosaic, and thus consider climb motion to be irrelevant for our system.

**Supplementary Figure 6.**
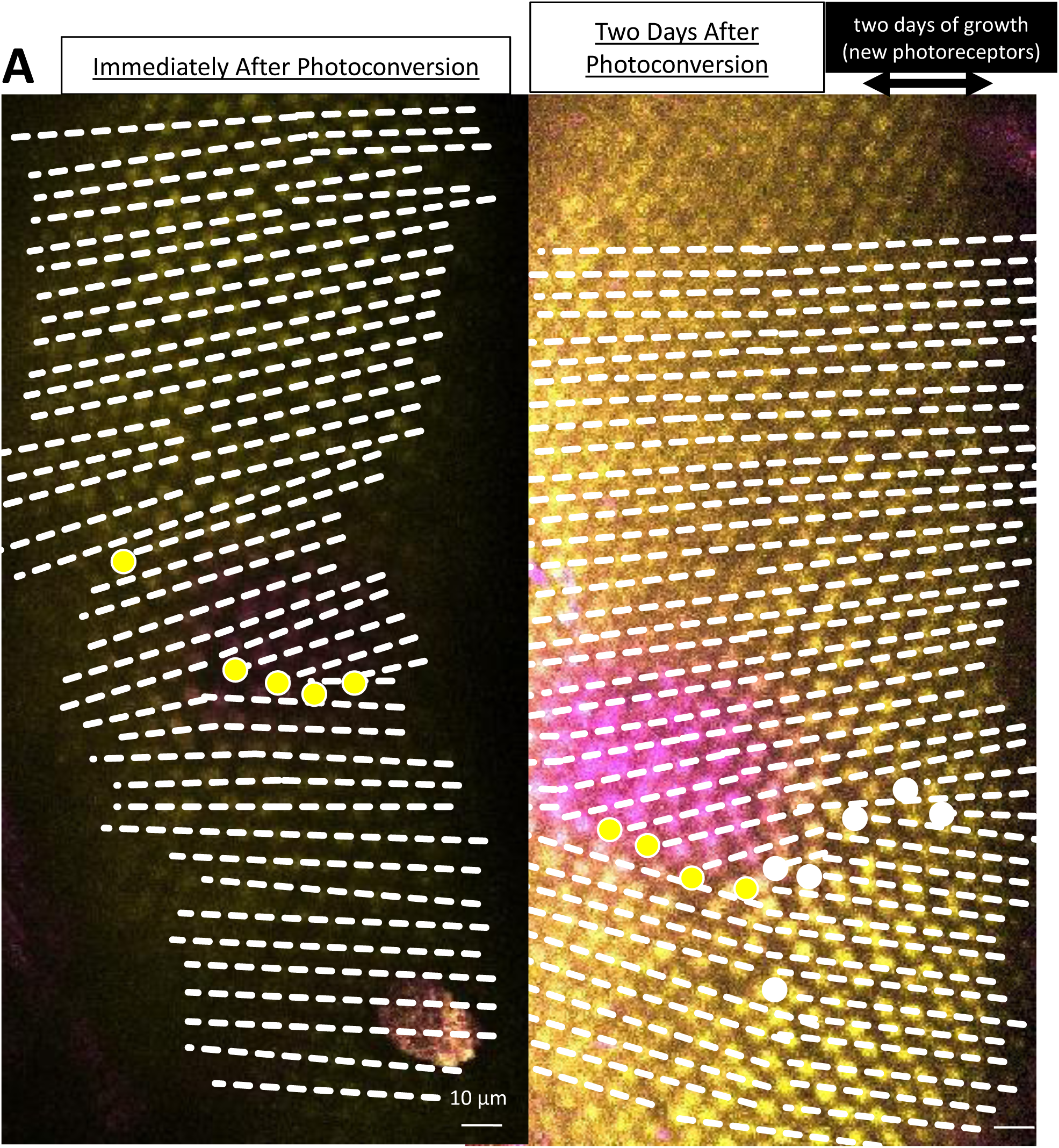
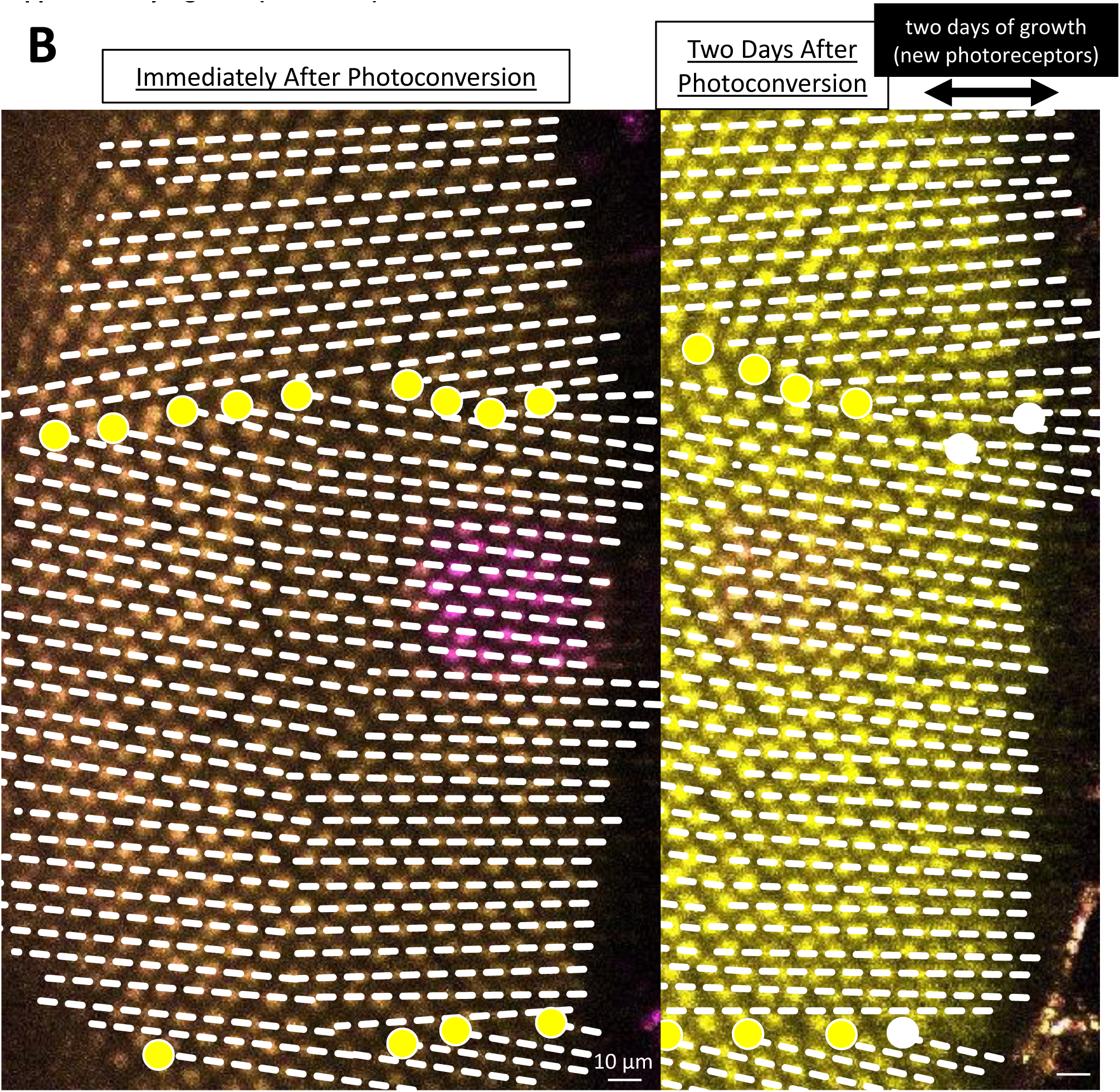
Further examples of photoconverted retinae in which we observe grain boundaries growing during initial cone mosaic formation. **A)** We trace rows of UV cones (white dashed lines) near Y-Junctions. The Y-Junctions observed immediately after photoconversion are indicated by yellow dots. The Y-Junctions observed in the newly incorporated after two days (newly incorporated) are indicated by white dots. The newly incorporated columns of UV cones are indicated by the double-sided black arrow. This is grain boundary 1 in Fig. 5E. B) All row tracing and Y-Junctions are denoted in the same way as in panel A. These are grain boundaries 4-1 and 4-2 in Fig. 5E.

**Supplementary Figure 7.**
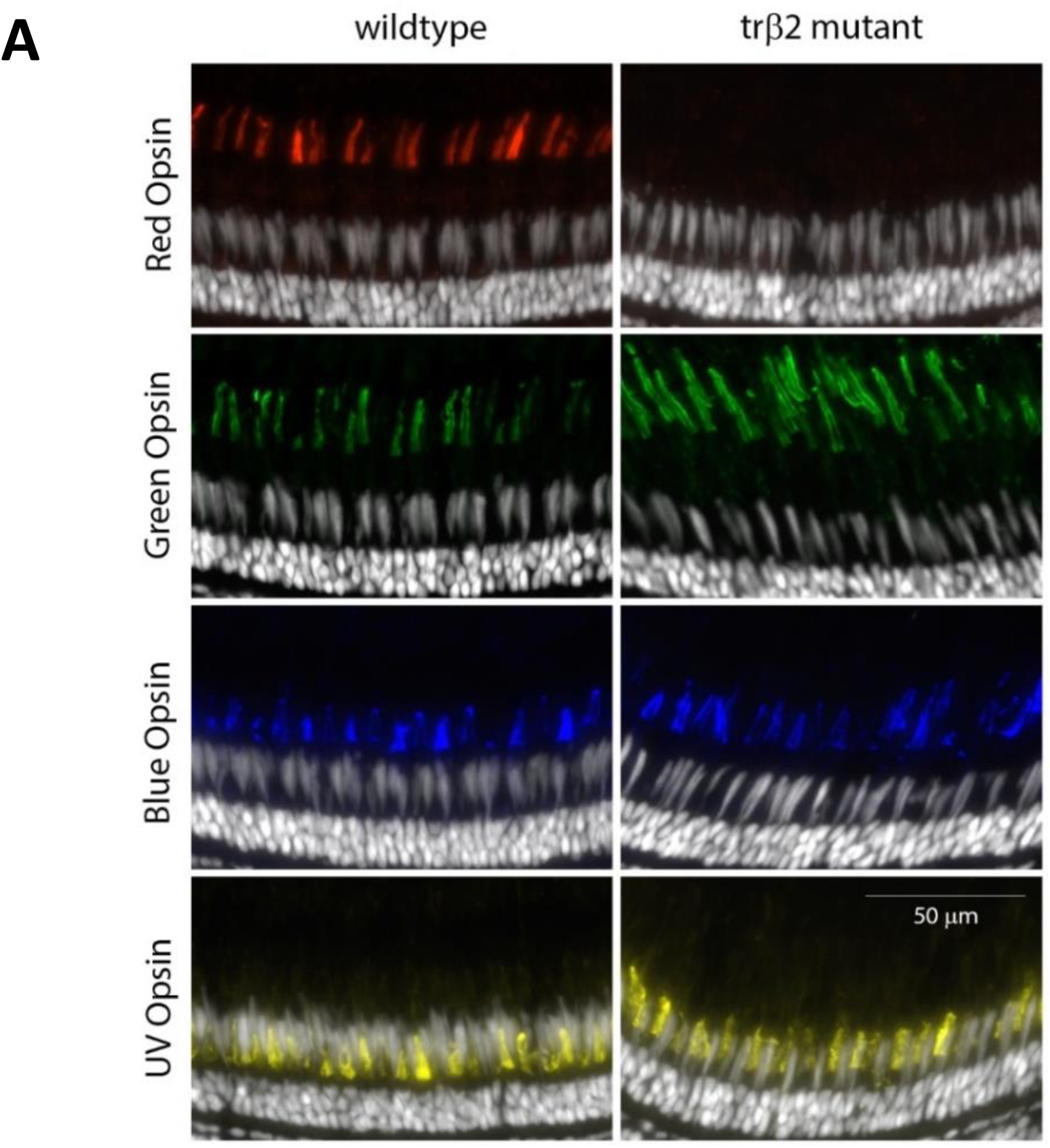
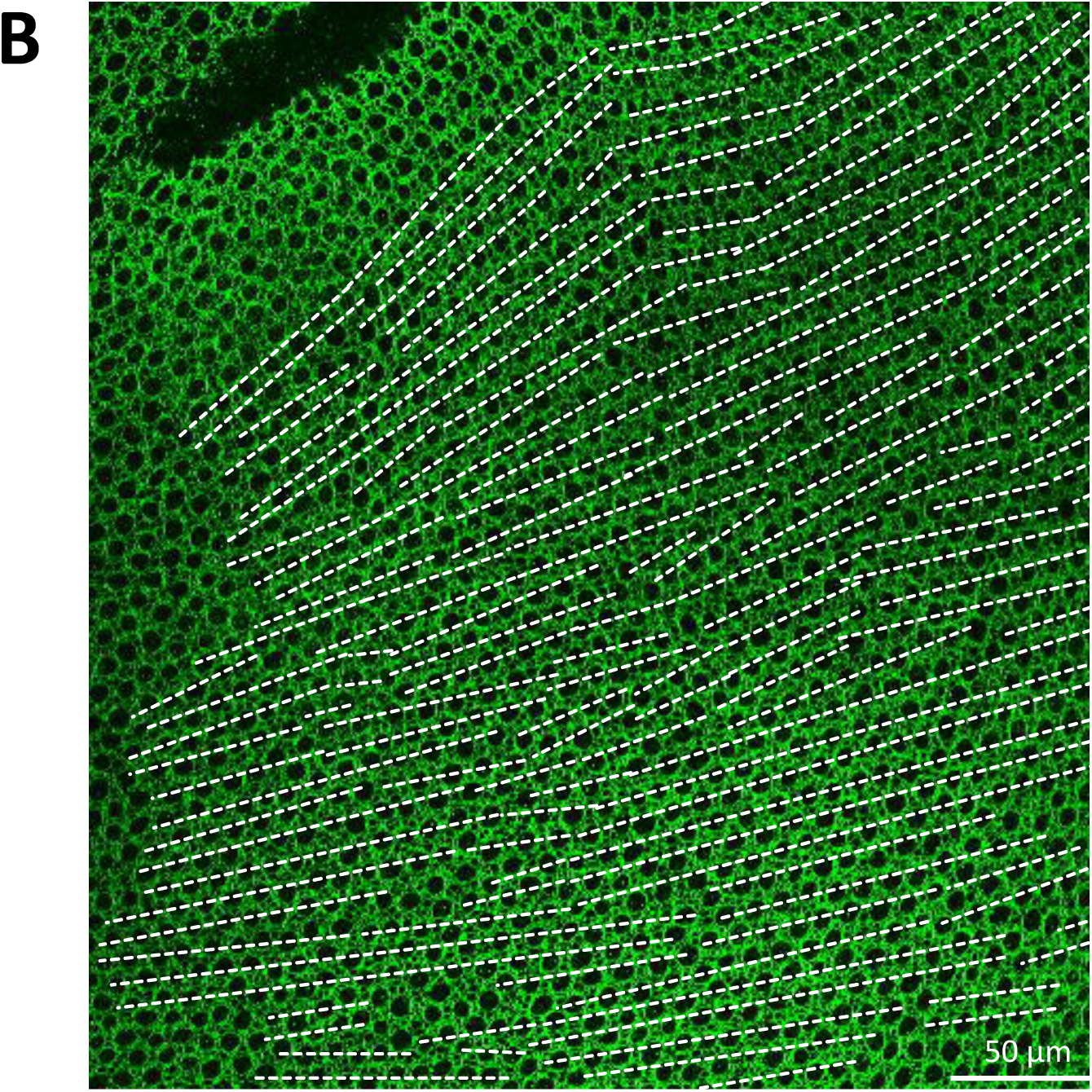
Mutation in the *trβ2* deletes Red cones, but not other cone subtypes. **A)** Immunocytochemistry for cone subtype specific opsins, Red opsin (red), Green opsin (green), blue opsin (blue), and UV opsin (yellow) in wildtype and *trβ2* mutant retinas. B) Flat-mount retinal preparation the *trβ2* mutant immunostained with ZO1 (green). The profiles of UV cone are large and rounded (see Fig. 6C-D). As a guide to the eye, we trace some rows of UV cones in the image with white dashed lines.

**Supplementary Figure 8.**
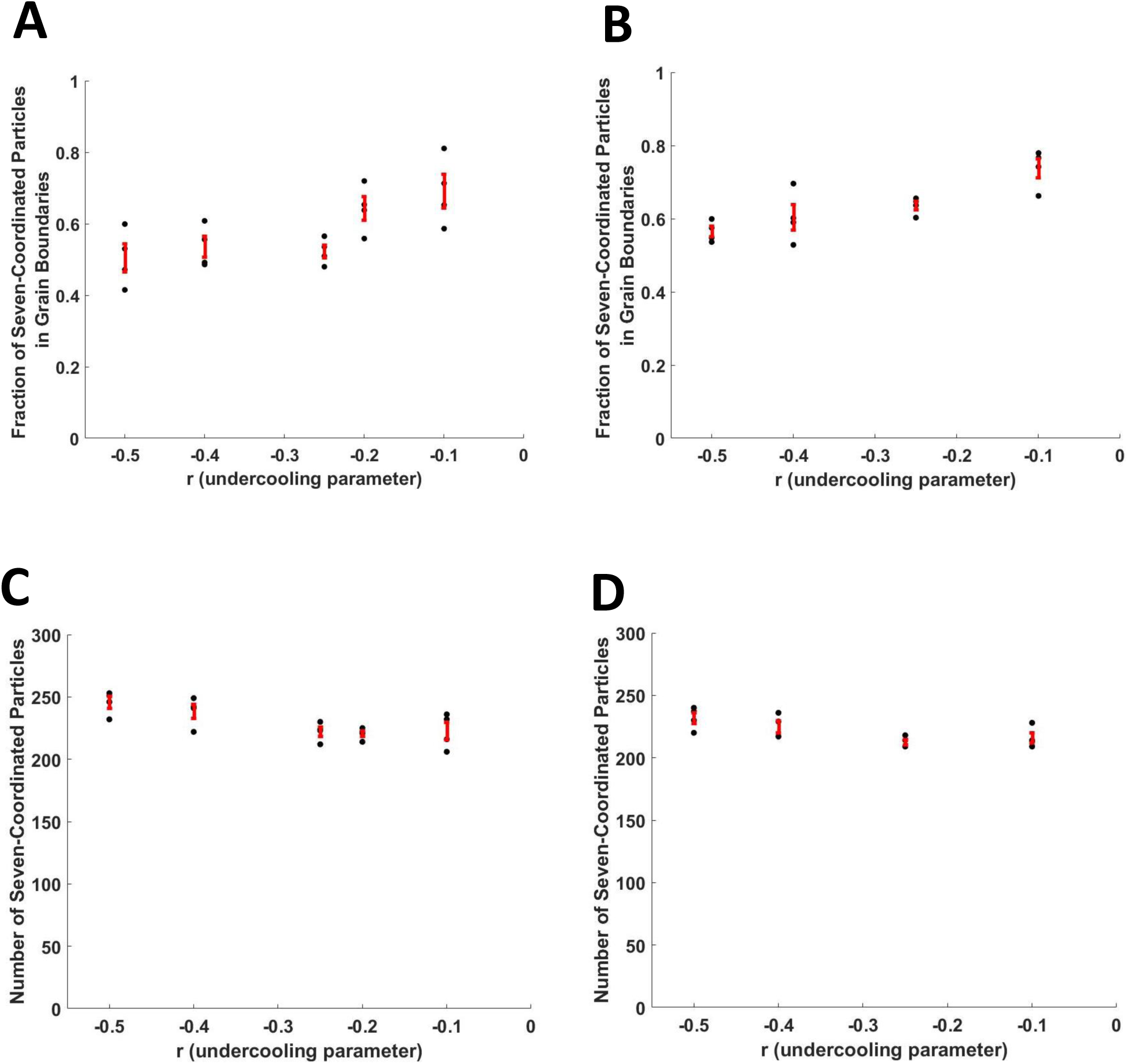
Scanning Parameters of Phase-Field Crystal Model. We take a one-dimensional cut of the two-dimensional phase diagram of the phase-field crystal model 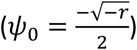 where *ψ*_0_ is the mean of the density modulation field and where *r* is the undercooling parameter. The number of initial rows on the cone frustum is two hundred. There are approximately ninety-five columns from the top of the cone frustum to the bottom. About two row insertions per added column are necessary to maintain constant cell-cell spacing on this cone. The degree of anisotropy is constrained by Fig. S2. A) The standard deviation of the white noise field, which is added to the first two columns, in these simulations is three-quarters. Along the one-dimensional cut of the PFC phase diagram, we measure the fraction of seven-coordinated particles in grain boundaries. B) The standard deviation of the white noise field in these simulations is one. Along the one-dimensional cut of the PFC phase diagram, we measure the fraction of seven-coordinated particles in grain boundaries. C) For the same simulations in panel A, we plot the number of seven-coordinated particles. D) For the same simulations in panel B, we plot the number of seven-coordinated particles.

**Supplementary Figure 9.**
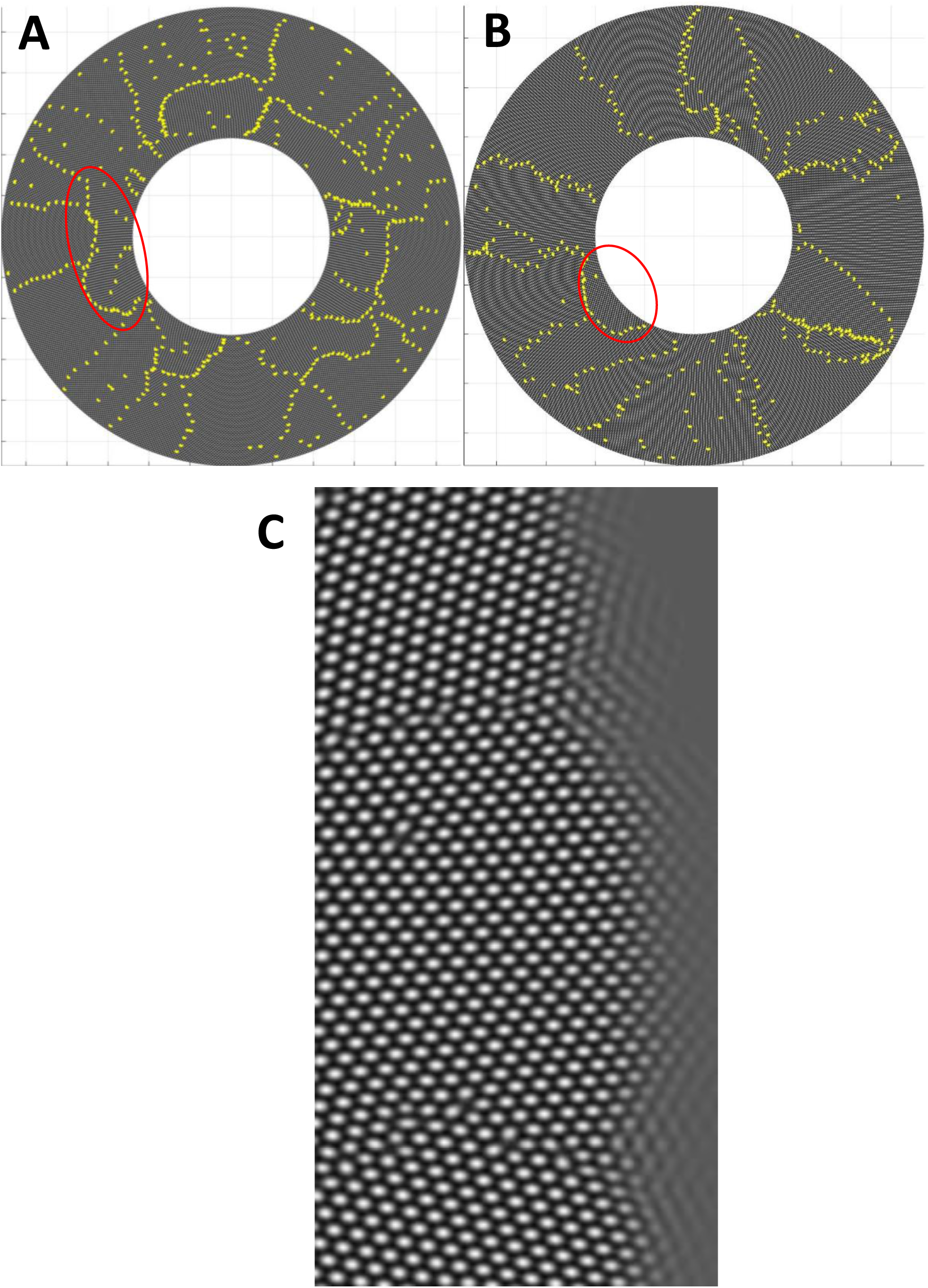
Additional insights generated by phase-field model of cone mosaic formation. **A)** Example of isotropic crystal growth (same example as Vid. 3) on cone frustum with an initial column as a prepattern. All seven-coordinated particles are denoted by yellow dots. Note the lines of seven-coordinated particles that do not radiate from the center of the cone to the periphery (example within red oval). These non-radiating lines of seven-coordinated particles are the result of a rotation of crystallographic orientation during growth of the isotropic crystal (*i.e.,* a domain rotation not observed in zebrafish retinae). B) Example of anisotropic crystal growth (same example as Vid. 5) on cone frustum with no initial column as a prepattern. With only white noise at the top of the cone in the initial conditions, the anisotropy of the crystal (*i.e.*, in the phase-field crystal free energy) selects the orientation and maintains that orientation during growth (in contrast with panel A). Even when a domain forms with the improper orientation (example within red oval), the domain rotates to the proper orientation during growth. C) Zoomed-in snapshot of an anisotropic phase-field crystal simulation on a cone. Note that near the grain boundaries (*i.e.*, where the domain rotation rotates), there is a lag in proper positioning of UV cones (*i.e.*, the density field remains poorly resolved) relative to growth of neighboring domains. This results in a characteristic V-shape.

**Supplementary Figure 10.**
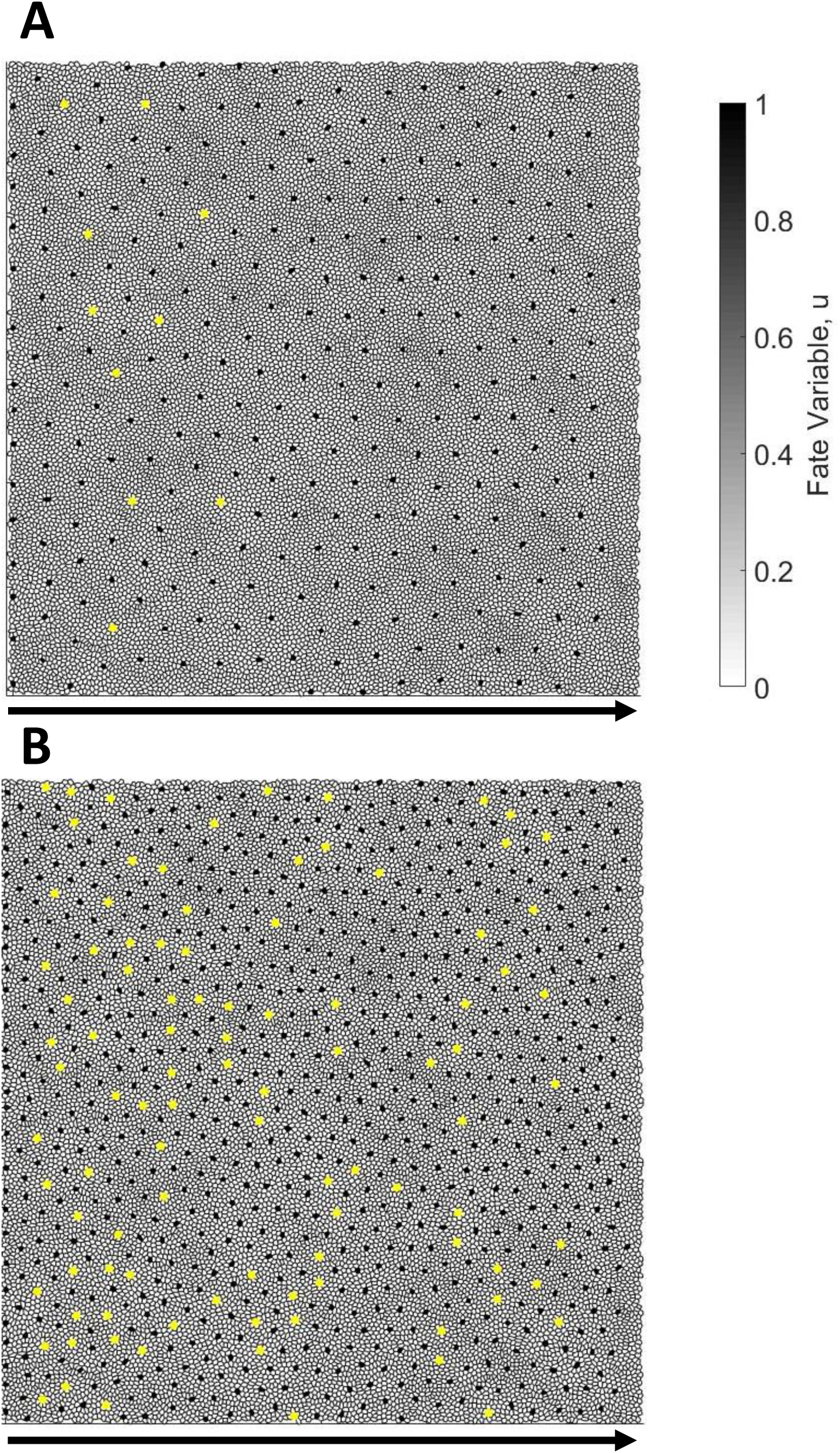
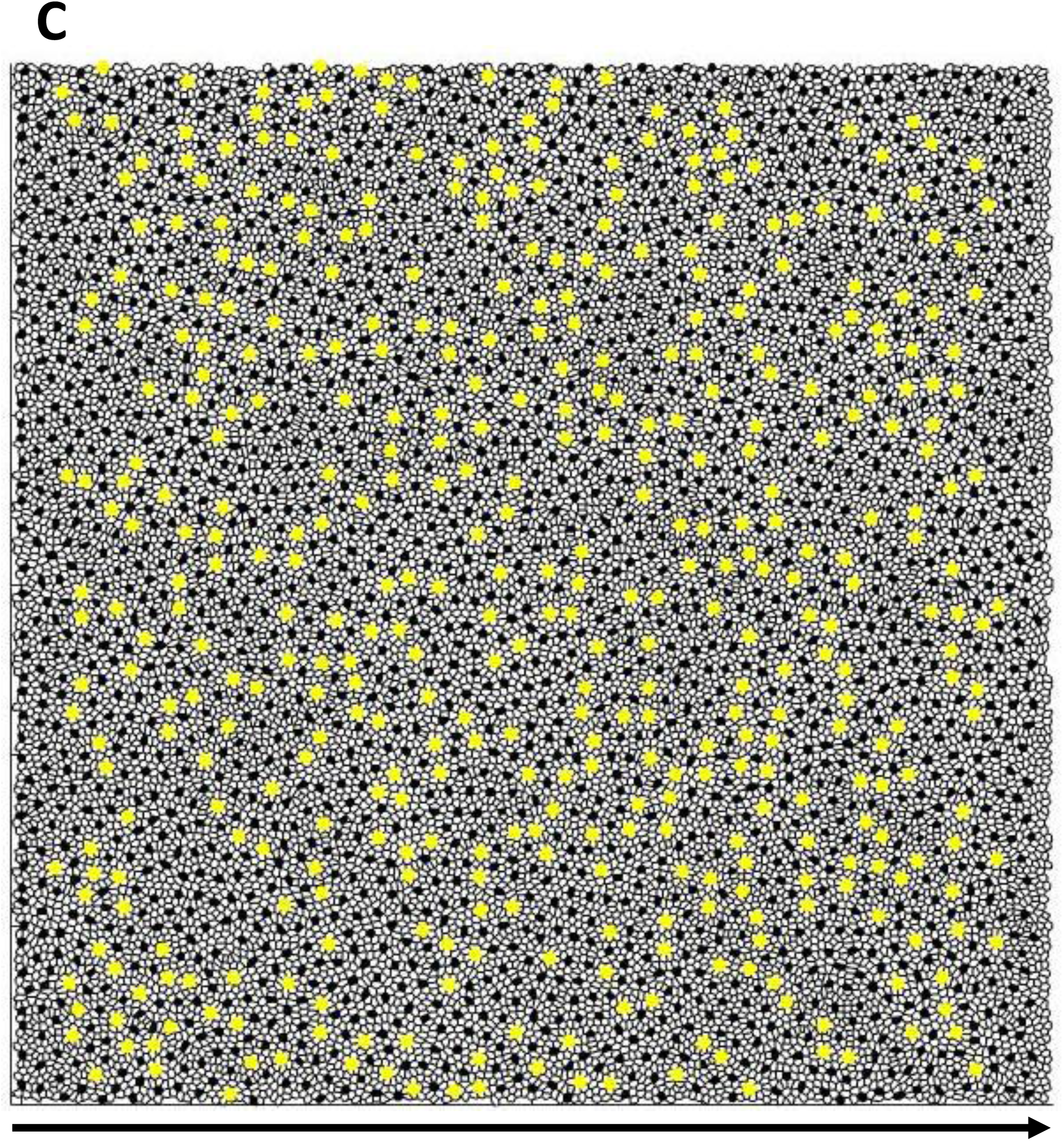
Lateral inhibition, with varying signaling ranges, in a disordered cell packing. A triangular lattice of *u* ≈ 1 cells forms on a square packing of 20000 cells with periodic boundary conditions. Defects (i.e., seven-coordinated) in this triangular lattice of *u* ≈ 1 cells are indicated by yellow dots. Initially, all cells are in the state (*u* = 0) because an external inhibiting signal is uniformly provided to all cells. Starting at *t* = 0, a wave of de-inhibition moves from left to right in the packing *l* (see Methods). The wave moves at a speed 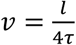 where *τ* is the time-scale of cell differentiation and *l* is the range of cell-cell signaling. In each panel, the black arrow denotes the direction of wave propagation. A) The signaling range is 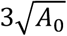, where *A*_0_is the mean cell area. This signaling range results in seven to eight *u* ≈ 0 cells between each pair of neighboring *u* ≈ 1 cells in the final pattern. Note that some defects are generated early in pattern formation (i.e., left side of packing), but the right side of the packing (i.e., more recently generated pattern) contains no defects. B) The signaling range is 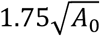, the same range as used in [28]. This results in about five *u* ≈ 0 cells between each pair of neighboring *u* ≈ 1 cells in the final pattern. Note that the entire packing contains defects. C) The signaling range is 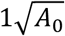, in the cone mosaic. This results in one to two *u* ≈ 0 cells between each pair of neighboring *u* ≈ 1 cells in the final pattern. Note that the entire packing contains defects. This image is enlarged relative to panels A-B for the sake of clarity.

## SUPPLEMENTARY TABLE CAPTIONS

**Supplementary Table 1.** Counts of reverse Y-Junctions. (i.e., row deletions) in regions of the retinae in which we traced rows. The same fish numbers are used in Fig. 3E. *There is a large-angle grain boundary in this retina, where patterning of the cone mosaic is slightly disrupted. There are potentially 10 additional reverse Y-Junctions associated with that large-angle grain boundary.

| Fish # | Number of (forward) Y-Junctions | Number of reverse Y-Junctions |
| --- | --- | --- |
| 1 | 155 | 0 |
| 2 | 166 | 5 |
| 3 | 221 | 0 |
| 4 | 275 | 5 |
| 5 | 249 | 14 |
| 6 | 184 | 2 |
| 7 | 182 | 10* |
| 8 | 285 | 7 |

**Supplementary Table 2.** Motion of Defects within Photoconverted Region. Here we quantify the motion of Y-Junctions within photoconverted regions. The fish labels are the same as in Figure 5E. Note that some fish are missing from this list (e.g., 4 and 8). These are fish which have grain boundaries in neighboring non-photoconverted regions, but do not have defects within the photoconverted region itself. For each fish, we fit the UV cone nuclei positions in the photoconverted regions to a plane in order to compute a triangulation. Because the photoconverted region is small relative to the radius of curvature of the retina, the UV cone nuclei positions are well fit (as quantified by RMSE) by a plane at both imaging times. In the UV cone triangulations, we check for bond flips near the defect core between photoconversion and later imaging (see Methods). If a Y-Junction glides by one unit, we denote that with a 1, and if a Y-Junction does not glide, we denote that with a 0.

| Fish Label | # of Y-Junctions in Photoconverted Region | Days between Photoconversion and Later Imaging | RMSE for Fit Plane at Day 0 ($\mu m$) | RMSE for Fit Plane at later imaging ($\mu m$) | Glide Motion of Defect(s) In Units of Rows |
| --- | --- | --- | --- | --- | --- |
| 1 | 2 | 2 | 0.85 | 0.63 | (0, 0) |
| 2 | 1 | 2 | 0.76 | 0.55 | 0 |
| 3 | 2 | 2 | 0.50 | 0.42 | (0, 0) |
| 5 | 1 | 2 | 0.70 | 0.26 | 1 |
| 6 | 1 | 2 | 1.0 | 0.59 | 0 |
| 7 | 1 | 2 | 0.48 | 0.63 | 1 |
| 9 | 1 | 2 | 0.96 | 0.53 | 0 |
| 10 | 2 | 3 | 1.1 | 0.62 | (0, 1) |
| 11 | 1 | 4 | 1.8 | 0.82 | 0 |
| 12 | 1 | 4 | 1.5 | 0.74 | 1 |
| 13 | 1 | 2 | 0.85 | 0.31 | 1 |
| 14 | 1 | 2 | 0.90 | 1.1 | 0 |
| 15 | 1 | 2 | 1.2 | 0.45 | 1 |

**Supplementary Table 3.** New Y-Junctions are incorporated preferentially near existing grain boundaries in live fish. The fish labels are the same as in Figure 5E and S6. We first list the number of Y-Junctions incorporated between photoconversion and later imaging (within the whole image, not just within the grain boundary). We list the average and median distances (i.e., along the margin axis) between a new Y-Junction and the nearest grain boundaries if new Y-Junction positions are uncorrelated with positions of existing grain boundaries. We, then, list the actual distances (i.e., along the margin axis) between newly incorporated Y-Junctions and the nearest existing grain boundary in the image. Fish 8 is omitted because, within the image, no new Y-Junctions are added between photoconversion and later imaging.

| Fish Label | # of Y-Junctions Added Between Photoconversion and Later Imaging | Mean Distance Between New Y-Junction and Nearest GB if Uncorrelated ($\mu m$) | Median Distance Between New Y-Junction and Nearest GB if Uncorrelated ($\mu m$) | Actual Distance Between new Y-Junctions and GB ($\mu m$) |
| --- | --- | --- | --- | --- |
| 1 | 6 | 84 | 73 | [9, 2, 15, 12, 12, 46] |
| 2 | 2 | 91 | 73 | [24, 4] |
| 3 | 3 | 87 | 73 | [1, 2, 65] |
| 4 | 4 | 45 | 44 | [24, 1, 5, 1] |
| 5 | 1 | 73 | 73 | 1 |
| 6 | 2 | 75 | 73 | [4, 1] |
| 7 | 3 | 74 | 73 | [11, 123, 42] |
| 9 | 2 | 73 | 73 | [18, 20] |
| 10 | 2 | 94 | 78 | [2, 2] |
| 11 | 8 | 43 | 39 | [12, 23, 45, 1, 0, 95, 20, 69] |
| 12 | 3 | 39 | 36 | [5, 11, 36] |

